# Ammonia leakage can underpin nitrogen-sharing among soil microbes

**DOI:** 10.1101/2024.04.08.588523

**Authors:** Luke Richards, Patrick Schäfer, Orkun S Soyer

## Abstract

Soil microbial communities host a large number of microbial species that support important ecological functions such as biogeochemical cycling and plant nutrition. The extent and stability of these functions are affected by inter-species interactions among soil microbes, yet the different mechanisms underpinning microbial interactions in the soil are not fully understood. Here, we study the extent of nutrient-based interactions among two model, plant-supporting soil microbes, the fungi *Serendipita indica* and the bacteria *Bacillus subtilis*. We find that *S. indica* is unable to grow with nitrate - a common nitrogen source in the soil - but this inability can be rescued, and growth restored in the presence of *B. subtilis*. We demonstrate that this effect is due to *B. subtilis* utilising nitrate and releasing ammonia, which can be used by *S. indica*. We refer to this type of mechanism as ammonia mediated nitrogen sharing (N-sharing). Using a mathematical model, we demonstrate that the pH dependent equilibrium between ammonia (NH_3_) and ammonium (NH_4_^+^) results in an inherent cellular leakiness, and that reduced amonnium uptake or assimilation rates can result in higher levels of leaked ammonia. In line with this model, a mutant *B. subtilis* - devoid of ammonia uptake - shows higher *S. indica* growth support in nitrate media. These findings highlight that ammonia based N-sharing can be a previously under-appreciated mechanism underpinning interaction among soil microbes and could be influenced by microbial or abiotic alteration of pH in microenvironments.

**Significance statement:** Soil microbial communities are an important factor in environmental nutrient cycling and subsequently plant health. *S. indica* is a well-studied plant growth promoting soil fungus but its inability to use nitrate, a major component of both agricultural/natural soils and crop fertilisers, may have important implications for agriculture and microbial ecology. We have demonstrated that *S. indica* is dependant on external sources of nitrogen in nitrate-only environments and these can be produced by *B. subtilis*, another common soil microbe. We then demonstrate that this nitrogen sharing interaction is likely mediated by leaked ammonia and that ammonia leakage is influenced by environmental pH. Ammonia leakage and sharing represent currently unexplored and potentially vital components of nutrient interactions between microbes in soil communities, with profound implications for microbiome community structure and subsequent consequences for soil biogeochemical cycling and crop health.

## 1 Introduction

The vast array of soil microorganisms and their interactions, the “soil microbial community”, confer a myriad of functions in soils. These functions underpin the contribution of soil community to biogeochemical cycles ([Condron et al., 2010], [Bhattacharjya et al., 2019], [Kuypers et al., 2018], [Kraemer, 2004]) and to plant growth and health ([Barea and Richardson, 2015], [Lemanceau et al., 2009], [Santoyo et al., 2017]). The latter contribution is mediated through microbial-processing of nutrients needed by the plant, suppression of plant pathogens, enhanced stress tolerance (reviewed by [Bakker et al., 2020]) and by modulation of plant hormone signalling ([Eichmann et al., 2021], [Osborne et al., 2023]). There is therefore a requirement to better understand soil microbial communities and the microbial interactions within, so to preserve and possibly increase crop/ soil fertility. Several studies have shown that abiotic environmental inputs influence soil and plant microbiome assembly ([Lagunas et al., 2023], [Lauber et al., 2009], [Gourmelon et al., 2016] and reviewed here [Santoyo et al., 2017]). At the community level, nutrient availability has been shown to strongly influence community diversity and it is suggested that high nutrient availability promotes the proliferation of few competitive species, while low nutrient availability promotes diversity ([Ratzke et al., 2020]). External nutrient supply, as well as micro-scale nutrient gradients, are therefore important factors in shifting the balance of metabolic microbial interactions, as shown in experiments with model systems ([Jiang et al., 2018], [Ratzke et al., 2020], [Kehe et al., 2021], [Kim and Or, 2017], [Fu et al., 2020], [Hoek et al., 2016]).

The environmental influences on community composition are likely mediated through direct effects on specific inter-microbial interactions, however, specific interactions among soil microbes are often not well-characterised. Generally speaking, interactions between microbes can take many forms from competitive to cooperative and can be mediated by a range of different mechanisms ([Cavaliere et al., 2017], [Coyte and Rakoff-Nahoum, 2019]). Among these, metabolic interactions through cross-feeding and auxotrophy are shown to be wide-spread in many different environments, including the soil ([Jiang et al., 2018], [Kazamia et al., 2012], [Ponomarova et al., 2017], [Uehling et al., 2019]). Auxotrophic interactions occur when one organism loses ability to synthesize a growth-essential compound, and instead receives this from another organism ([D’Souza and Kost, 2016]). Among specific soil microbes, previous work has highlighted mutual growth promotion between the fungus *Mortierella elongata* and bacterium *Burkholderia* BT03, mediated in-part by fungal organic acids ([Uehling et al., 2019]). An auxotrophic interaction has also previously been identified, via vitamin B1 (thiamine), between the bacterium *Bacillus subtilis* and the fungus *Serendipita indica* ([Jiang et al., 2018]). This specific interaction is notable, since *S. indica* is an important soil fungi believed to promote plant growth and stress tolerance across a range of plant species ([Qiang et al., 2012], [Varma et al., 1999], [Weiß et al., 2016]). The inability of *S. indica* to produce thiamine means that its beneficial activity towards plants is dependent on nutrient input from other soil organisms such as *B. subtilis* or the plants themselves.

In addition to its inability to produce thiamine, *S. indica* has been shown previously to have impaired growth in media with nitrate as the only N-source, and lacks genes coding for nitrate transporters and, nitrate and nitrite reductases ([Zuccaro et al., 2011]). Considering that nitrate is a major nitrogen source in global soils and agricultural practices ([Hill et al., 2011], [Lagunas et al., 2023], [Craswell, 2021], [Walvoord et al., 2003], [Rao and Puttanna, 2000]), this raises a question about why fungi such as *S. indica* would lose there ability to assimilate such an abundant nitrogen source and how they sustain themselves in nitrate-dominant environments. One possible answer to these questions could be that alternative nitrogen sources that are easier to utilise to be readily co-existing with nitrate, in nitrate-rich environments due to activities of nitrate utilising bacteria.

Here, we have explored this hypothesis using the *B. subtilis* – *S. indica* pair as a model bacteria – fungi interaction system. We reconfirm the incapability of *S. indica* to use nitrate as a sole nitrogen source and demonstrate that this incapacity is ameliorated by the presence of *B. subtilis*. We show that this effect is mainly due to ammonia, which we find to be released into the media by *B. subtilis*. This finding is supported by a mathematical model, showing that ammonia can be readily leaked by cells due to its pH dependent equilibrium with ammonium and the high permeability of the latter through cell membranes. Thus, our results show that inevitable leakage of ammonia can act as a nitrogen sharing mechanism among soil microbes. This highlights the possible importance of incidental leakage of membrane-permeable metabolites in the development of auxotrophic interactions and the maintenance of soil microbial community stability.

## 2 Results

### 2.1 S.indica lacks the ability to assimilate nitrate

Both bacteria and fungi, generally, use the same metabolic pathways for the assimilation of nitrogen into amino acids (Figure 1) ([Crawford and Arst Jr, 1993], [Lin and Stewart, 1997]). Nitrate can be taken up by active transport, via nitrate transporters (NTR), and reduced to nitrite and then ammonium by nitrate reductases (NR) and nitrite reductases (NiR). Ammonium can then be assimilated by two enzyme-facilitated reactions: glutamate dehydrogenase (GDH) catalyzes the NADH-dependent reversible reaction of *α*-ketoglutarate with ammonium, resulting in glutamate, while glutamine synthase (GS) catalyzes the reaction of ammonium with glutamate, resulting in glutamine. A third enzyme, glutamate synthase (GOGAT), catalyzes the re-generation of glutamate by a reaction that combines glutamine and *α*-ketoglutarate (Figure 1). Using the pBLAST tool ([Altschul et al., 1997]), and genes from *Bacillus subtilis* as a reference, we searched both the *S. indica* and *Serendipita vermifera* NCBI sequence repositories for GDH, GS, and GOGAT homologs, as well as for NTR, NR and NiR (see *Methods*). For GDH, GS and GOGAT we found likely homologs in both fungal species with a *>*25% sequence identity match over *>*89% of the query sequence length. For NTR, NR and NiR possible homologs were found in the *S. vermifera* genome with *>*26% sequence identity match over *>*42% of the query sequence length but no hits were identified in *S. indica* for NTR and those identified for NiR and NR had lower identity and length than that quoted for *S. vermifera*. Additionally the annotations for those hits do not match NRs and NiRs (Data file S1). To explore this further, we used the *Ogatea angusta* formerly *Hansenula polymorpha* yeast species’ nitrate assimilation genes (OaYNT1 nitrate transporter, OaYNR1 nitrate reductase and OaYNI1 nitrite reductase [ÁVILA et al., 1998]) to search the *S. indica* and *S. vermifera* genomes for homologous genes. This represents an example of a well-characterised nitrate-assimilating fungus ([Siverio, 2002]). Likely homologs were identified for all three genes in the *S. vermifera* genome while only the nitrate reductase (OaYNR1) produced hits in the *S. indica* genome (Data file S1). These had low sequence similarity over only the latter portion of the query sequence where the NADH binding domain is located in other fungal NRs ([Campbell and Kinghorn, 1990]). These results, and in particular the lack of NTRs, suggest that *S. indica* will be unable to use nitrate as the sole nitrogen source.

**Figure 1:**
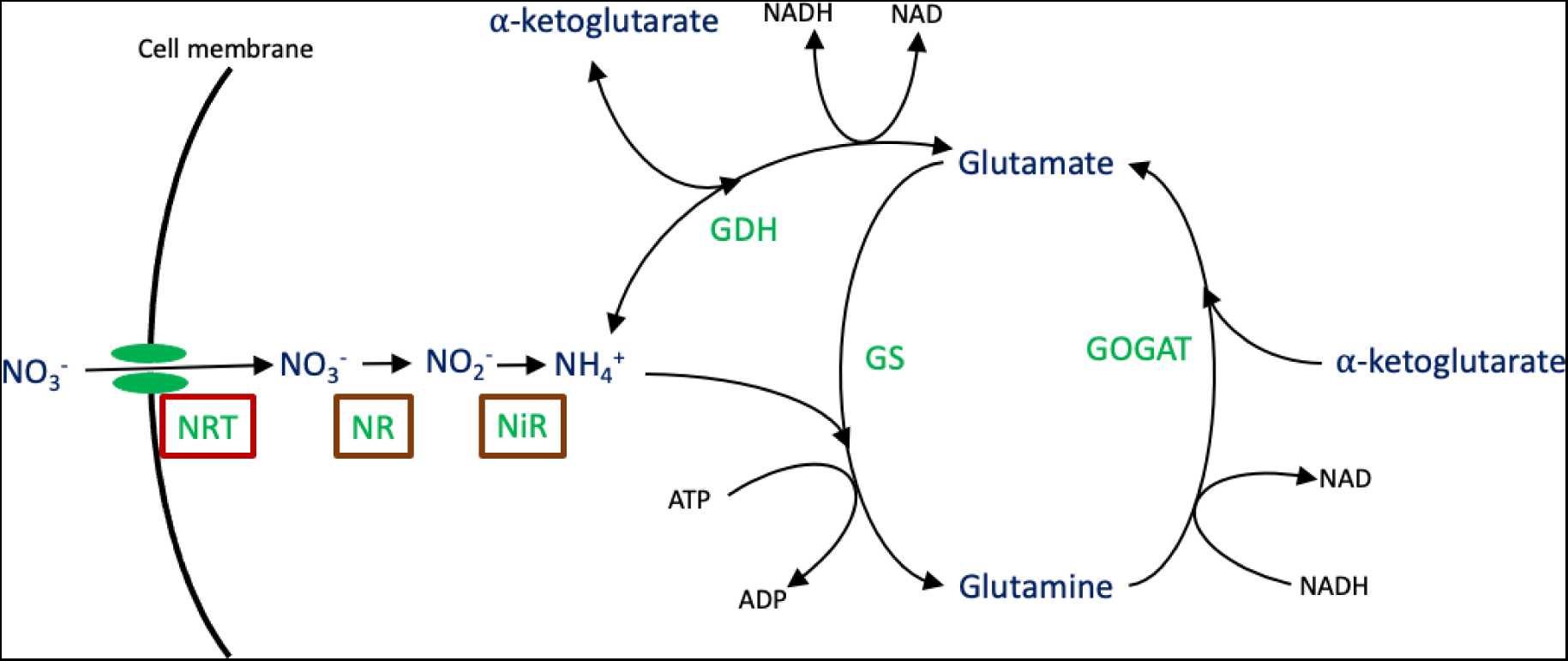
General pathway of nitrogen assimilation. Major metabolites are indicated in dark blue, enzymes in green and boxed enzymes are those not found in *S. indica*.

To experimentally test the capability of *S. indica* to use nitrate, we grew *S. indica* spores on ATS media containing different nitrogen sources: Nitrate, ammonium and glutamine (see *Methods*). *S. indica* displayed extremely reduced growth on media containing nitrate when compared with ammonium and glutamine (Figure 2A). It cannot be ruled out that such low growth is mediated solely by nitrogen reserves stored in spores. Supporting this possibility, quantification of growth in liquid culture through dry-weight measurements showed similar level of growth in nitrate only media compared to media with no nitrogen source (Figure S1). Growth in glutamine-containing media, which could be seen as a mimic of the availability of amino acids/ organic nitrogen in soil, was comparable to that with ammonium (Figure 2A). These results, together with genetic information given above, show that *S. indica* is unable to assimilate nitrate.

**Figure 2:**
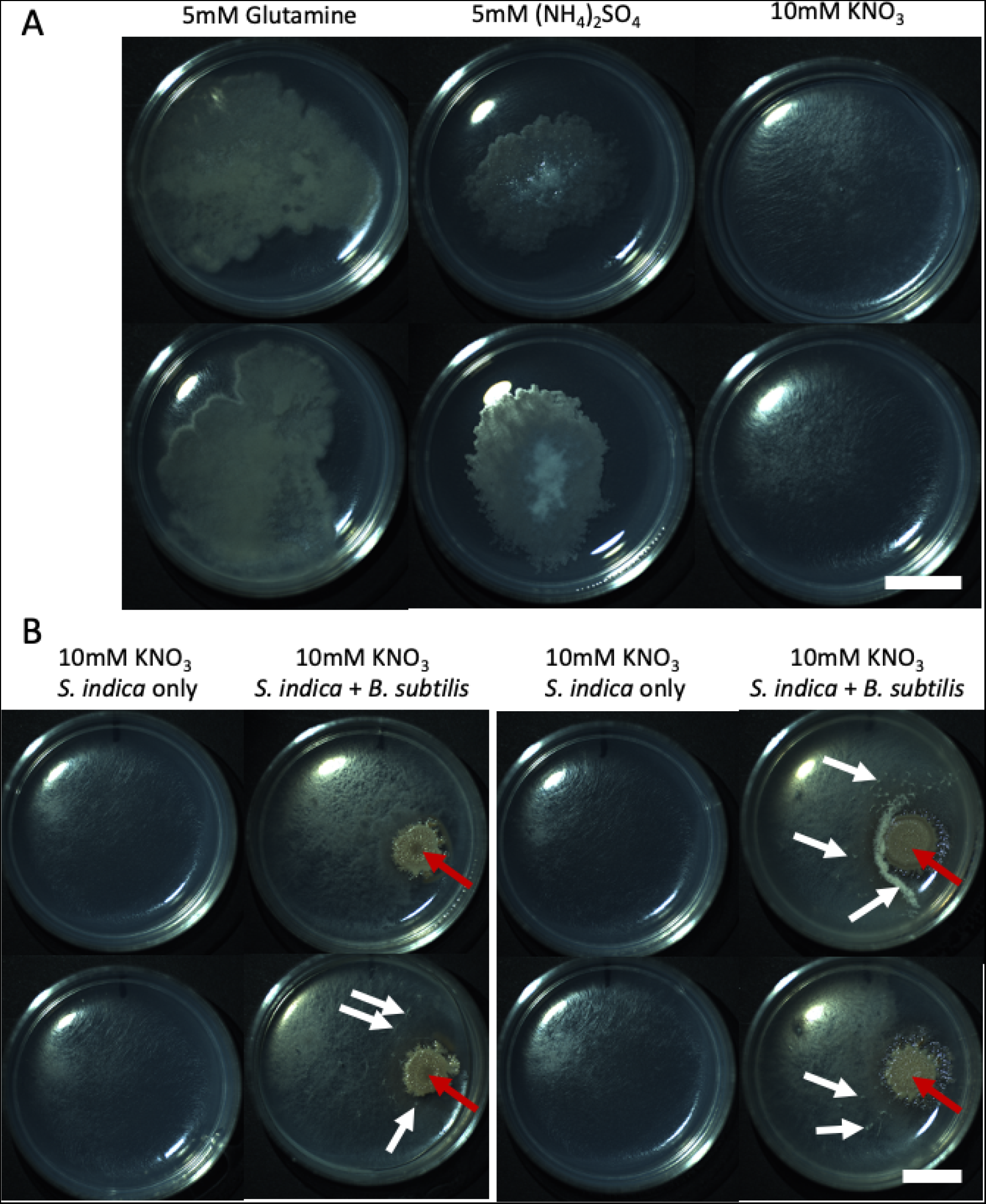
A) *S. indica* growth after 42 days on ATS media supplemented with the indicated N-source. B) *S. indica* growth after 42 days in isolation or in the presence of *B. subtilis* on ATS media supplemented with 10mM KNO_3_. *B. subtilis* inoculum was added 2 days after *S. indica* inoculation. On co-culture plates, large *B. subtilis* colonies are visible on the right, indicated with red arrows. In co-culture the mat of *S. indica* mycelia appears generally more dense but is also accompanied by “fluffy” protrusions from the media, indicated with white arrows. The two sets of images shown on the left and right panels are from replicate experiments. Scale bar is 1cm and applies to all panels.

### 2.2 S. indica growth in nitrate media is significantly enhanced in the presence of B. subtilis

Given that *S. indica* can readily use ammonia and glutamine, we hypothesised that soil bacteria may be able to provide *S. indica* with these alternative nitrogen sources in a nitrate-dominated environment. To investigate this, *S. indica* was grown on ATS-nitrate plates with and without the addition of *B. subtilis* (see *Methods*). The presence of *B. subtilis* improves the growth of *S. indica* on agar plates (Figure 2B), leading to the hypothesis that *B. subtilis* is providing some exuded nitrogen compound capable of diffusing across the media and being taken up by *S. indica*.

To further investigate this possibility, we assessed growth of *S. indica* in liquid culture in the presence of *B. subtilis* supernatant. Trial experiments indicated that, qualitatively, *B. subtilis* supernatant (collected over a range of optical densities) was able to promote the growth of *S. indica* in liquid culture (Figure S2A). Nitrate and ammonium quantification of the *B. subtilis* supernatant samples used in these experiments shows consumption of nitrate as OD increase and a production of ammonium (Figure S2). At higher optical densities the growth promotion was reduced, possibly because of a re-consumption of ammonium by *B. subtilis* at high bacterial cell densities (Figure S2). To re-confirm this growth-promotion result quantitatively, we designed experiments focusing on *B. subtilis* supernatant collected at an OD below 2 (Figure S3 and *Methods*). The *B. subtilis* supernatant addition greatly increased the dry mass of *S. indica* grown in liquid culture over 1 week (Figure 3A), supporting the hypothesis that exudates from *B. subtilis* are used as a nitrogen source by *S. indica*. To further confirm this observation, we repeated the supernatant supplementation experiment presented in Figure 3A and again found a significant growth promotion of *S. indica* with *B. subtilis supernatant*(Figure S4). We also quantified the nitrate in supernatants from these experiments (see *Methods*)and were able to show a clear reduction in the amount of nitrate in *B. subtilis* supernatants which was not seen in the supernatants of *S. indica* (Figure S5). Thus, *B. subtilis* growth results in nitrate consumption and a release of a nitrogen source into supernatant that can be used by *S. indica*.

**Figure 3:**
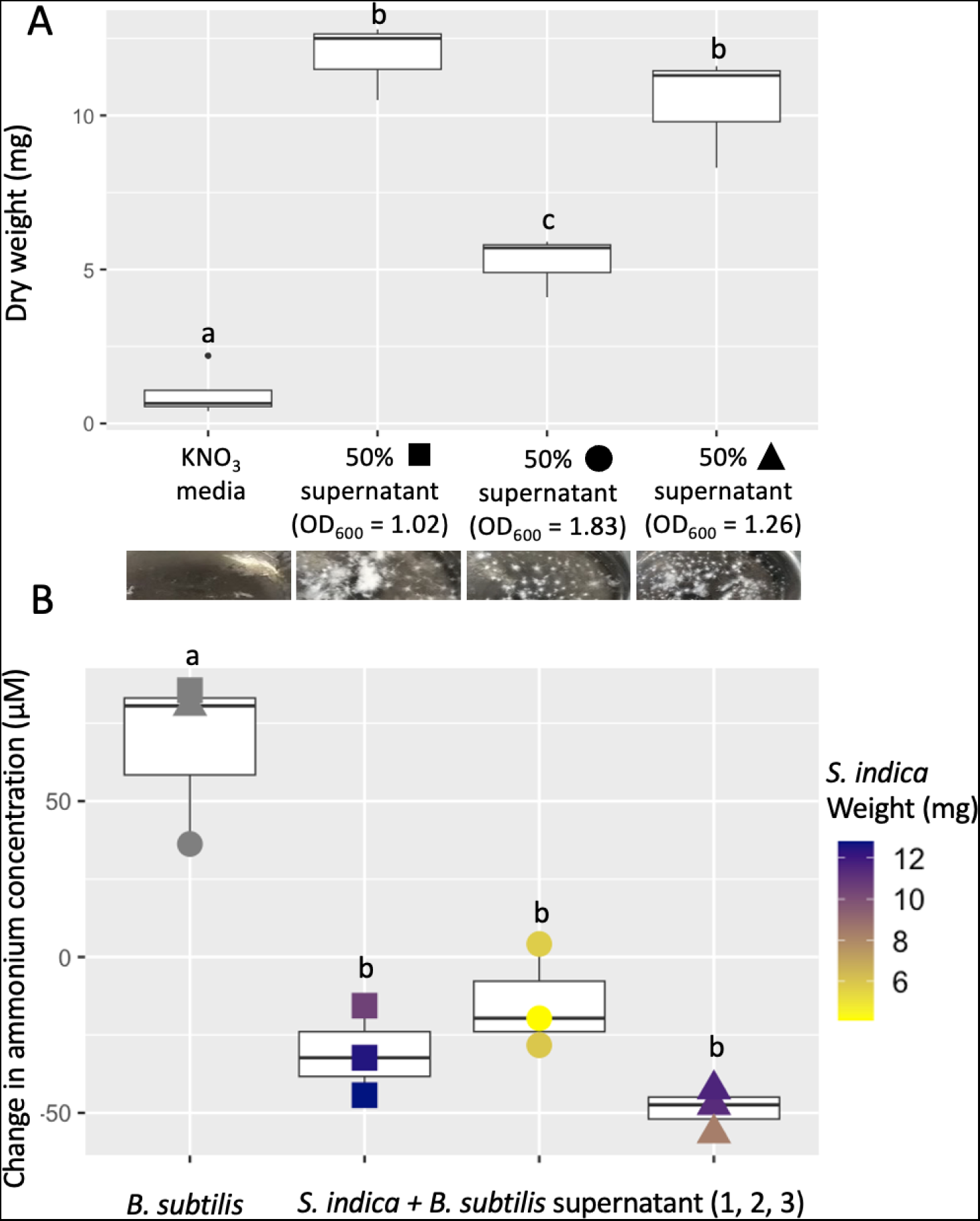
A) Dry weight of *S.indica* growth after 1 week of liquid culture in ATS media supplemented with 10mM KNO_3_ or a 50:50 mixture of this media and supernatant of a *B. subtilis* culture grown in the same media. Harvesting OD_600_ for this supernatant is indicated on the x-axis. B) Difference between end-point and starting NH_4_^+^ concentration, as measured by HPLC, in *S. indica* and *B. subtilis* cultures in conditions indicated on the x-axis. Where applicable *S. indica* dry weight (from A) for each individual point is indicated with a colour scale. For both figures mid point indicates median, edges of boxes indicate lower and upper quartiles and ends of whiskers indicate maxima and minima. Different letters above boxes indicate a significant difference (TukeyHSD p *<* 0.05). Plot point shapes indicate the OD_600_ of the *B. subtilis* culture at the time of supernatant harvest (from A): diamond - NA, square - 1.02, circle - 1.83, triangle - 1.26.

### 2.3 *S. indica* uses ammonia released by *B. subtilis* to grow in the nitrate-only media

To identify the compounds in the supernatant of nitrate-grown *B. subtilis* that *S. indica* could use as N-source, we analysed supernatant samples for different nitrogen compounds by HPLC (see *Methods*). We could not detect any free amino acids in any of the supernatant samples other than those collected at OD greater than 4 (Data file S1). We also tested supernatant samples for the presence of protein-derived amino acids by subjecting them to acid hydrolysis prior to HPLC quantification. The sum of protein-derived amino acids was not significantly different between samples and generally the amounts detected were low (Figure S5). High amounts of ammonia and taurine were detected in all of these hydrolysis-treated samples, including media only controls, indicating that hydrolysis can result in production of these compounds from nitrate.

The lack of any significant excreted or protein-derived amino acids in supernatant of nitrate-grown *B. subtilis*, led us to hypothesise that the effect of the supernatant on *S. indica* could be due to ammonium. Indeed, we found low but significant levels of ammonium in all *B. subtilis* supernatants prior to hydrolysis (Figure S6). This suggests that *B.subtilis* grown in nitrate releases ammonium into the environment and that this ammonium is consumed by *S. indica* when grown with *B. subtilis* supernatant. Surprisingly, we also found production of ammonium, albeit less, in *S. indica* supernatants; both with nitrate-only and with supernatant-ameliorated media (Figure S6). The former finding indicates that spores have some stored nitrogen source, in line with the observation that very limited growth is possible in media with no nitrogen (Figure S1). The finding that ammonium is present, in varying amounts, in all supernatants leads us to ask if there is an inevitable leakage of ammonium. Indeed, it has been noted that ammonia (NH_3_) exists in equilibrium with ammonium (NH_4_^+^) in the media and inside the cell, and that the former can readily leak out of the cell due to its high permeability to the membrane ([Kim et al., 2012]). If leakage of ammonia is an unavoidable process, this would also explain the observation of ammonium in *S. indica* only samples, as some of the nitrogen stored in spores would be lost through ammonia leakage during its utilisation. Leakage of ammonia would also complicate the assessment of ammonium usage by *S. indica* when grown in *B. subtilis* supernatant, as consumption and leakage would occur simultaneously. In order to overcome this difficulty and best represent ammonium consumption by *S. indica* in the presence of *B. subtilis* supernatant we corrected the measured ammonium values by subtracting the mean ammonium concentration observed in *S. indica* only cultures (Figure S6) as an expected value of ammonium leakage from nitrogen stored in spores. Using these corrected values, we then calculate the change in ammonium concentration over the course of the experiment and compared these to the change in ammonium concentration in *B. subtilis* only cultures (Figure 3). We find that in cultures where *S. indica* is placed in media with *B. subtilis* supernatant there is clear consumption of ammonium. This effect is correlated with *S. indica* growth (Figure 3), supporting the conclusion that *S. indica* is using this leaked ammonium to support growth.

To further verify that ammonia levels found in *B. subtilis* supernatant can promote *S. indica* growth to the levels seen, we supplemented *S. indica* with the same levels of ammonium found in *B. subtilis* supernatant. We found that a supplementation of ammonium, equivalent to 37.5µM nitrogen can significantly enhance *S. indica* growth in nitrate media (Figure S7). Further confirming ammonia as mediater of the *B. subtilis* and *S. indica* interaction in these experiments, the lowest ammonium producing *B. subtilis* culture - one with the highest OD_600_ - produced the smallest growth benefit to *S. indica* (Figure 3). These findings strongly support the notion that the growth promotion effect of *B. subtilis* supernatant on *S. indica* in the nitrate-media is due to ammonium transferred between these organisms.

### 2.4 A simplified mathematical model of ammonia assimilation highlights the role of environmental pH on ammonia leakage and microbial interactions

To contextualise the above experimental results, we explored a simplified cellular growth model involving nitrate assimilation and ammonia leakage from cells. Dissolved gaseous ammonia (NH_3_) exists in an equilibrium with ammonium (NH_4_^+^), where the former readily diffuses rapidly across cell membranes whereas the latter must be actively transported ([Kim et al., 2012]).

Building on from a previously published model of ammonia assimilation ([Kim et al., 2012]), we first implemented a differential equation model with two compartments, representing a cell population growing in a homogenous environment (see *Methods*). We considered a given internal ammonium production rate, mimicking nitrate uptake and subsequent reduction (Figure 4A). Given this model, we explored the effect of key parameters, such as active ammonium uptake rate, ammonium assimilation rate (i.e. biomass incorporation rate), as well as the environmental pH level. We found that increasing uptake or assimilation rates reduces loss of ammonia to the media, although a certain level of leakage is always present even at high uptake and assimilation rates (Figure 4B&C). As can be expected, lowering the uptake or assimilation rate increases the extent of leakage (Figure 4B&C). This effect can be influenced by the environmental pH, where the impact of lowering the uptake or assimilation rate increases with lower environmental pH (Figure 4B&C). In other words, the ammonia leakage (for a given uptake and assimilation rate) becomes worse with decreasing environmental pH. The above analysis suggests that organisms must maintain a high ammonium uptake and assimilation rate to negate the effects of ammonia leakage, and do so more extensively under low pH conditions. It may also suggest that that it can be beneficial for “ammonium-scavenging” organisms to reduce their external pH, so to induce leakage in others, provided they can achieve higher uptake and assimilation rates.

**Figure 4:**
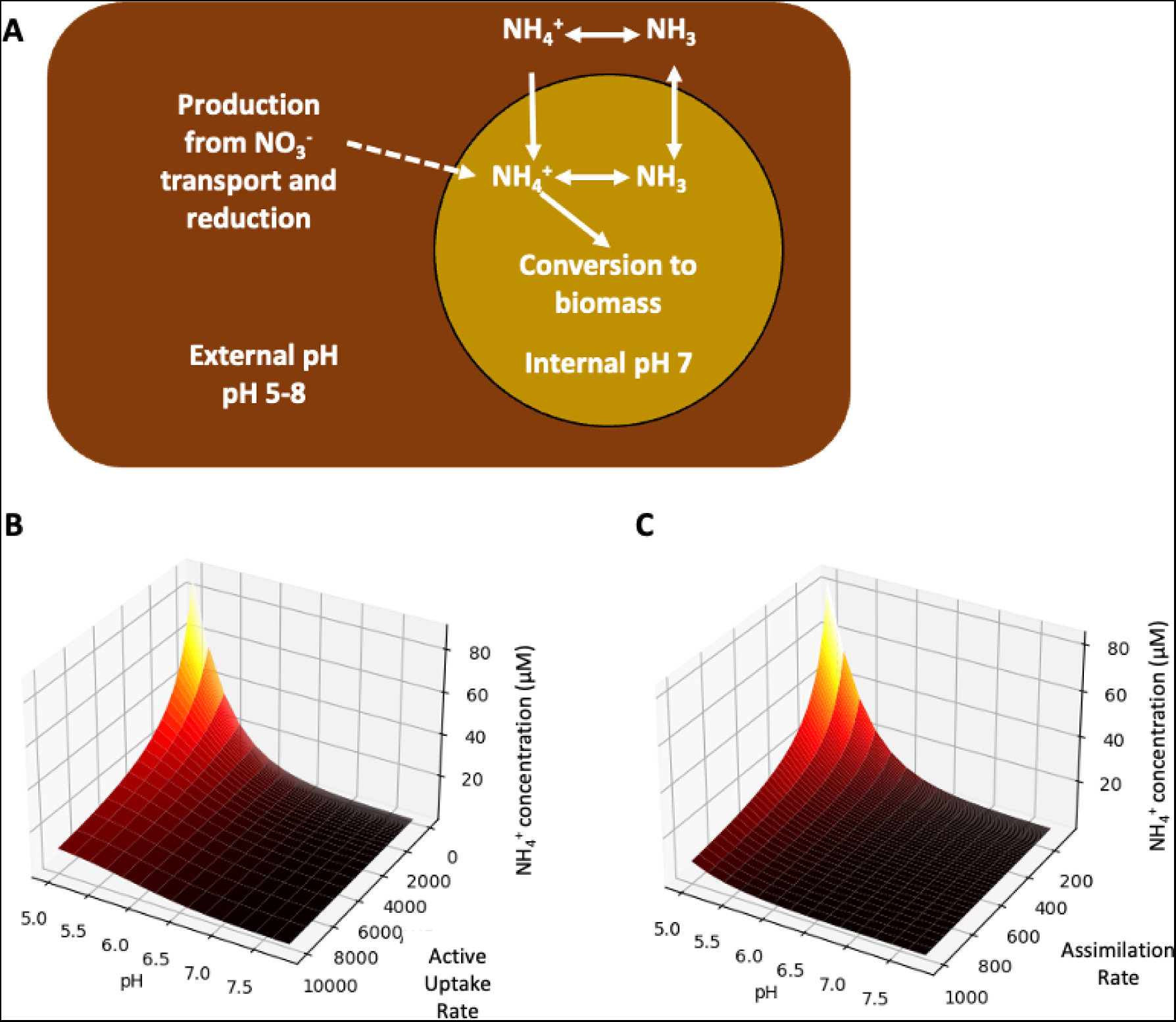
A) Diagram representing flow of N between compartments considered in our model and the pH values considered for each compartment. B&C) Model simulations depicting the external NH_4_^+^ concentration given various pH and B) Active uptake rates or C) Assimilation rates for a single cell with a constant rate of internal NH_4_^+^ production.

### 2.5 Reduced ammonia uptake in *B. subtilis* results in higher pH impact on its growth and increases its ability to support *S. indica* growth in nitrate media

One key consideration arising from the model is that the reduction of active uptake of ammonium by a given cell would increase its leakage of ammonia to the environment, and that this effect would be further enhanced under lower pH. To test this theoretical finding, we used an ammonium uptake mutant (*B. subtilis* 168ΔamtB) and its wild-type counterpart and assessed their ability to grow under different pH environments. These experiments were performed in homogenous environment to mimic the situation modeled in Figure 4A (see *Methods*). Growth of wild-type *B. subtilis* strain is not affected significantly by a pH change from 6.8 to 5.8, whereas the mutant strain with reduced active uptake of ammonium displays significantly lower growth under reduced pH (Figure 5). This supports the theoretical prediction that cells with reduced ammonium uptake will have a pH dependent impact on their ability to keep ammonium and therefore might be affected in their growth rate.

**Figure 5:**
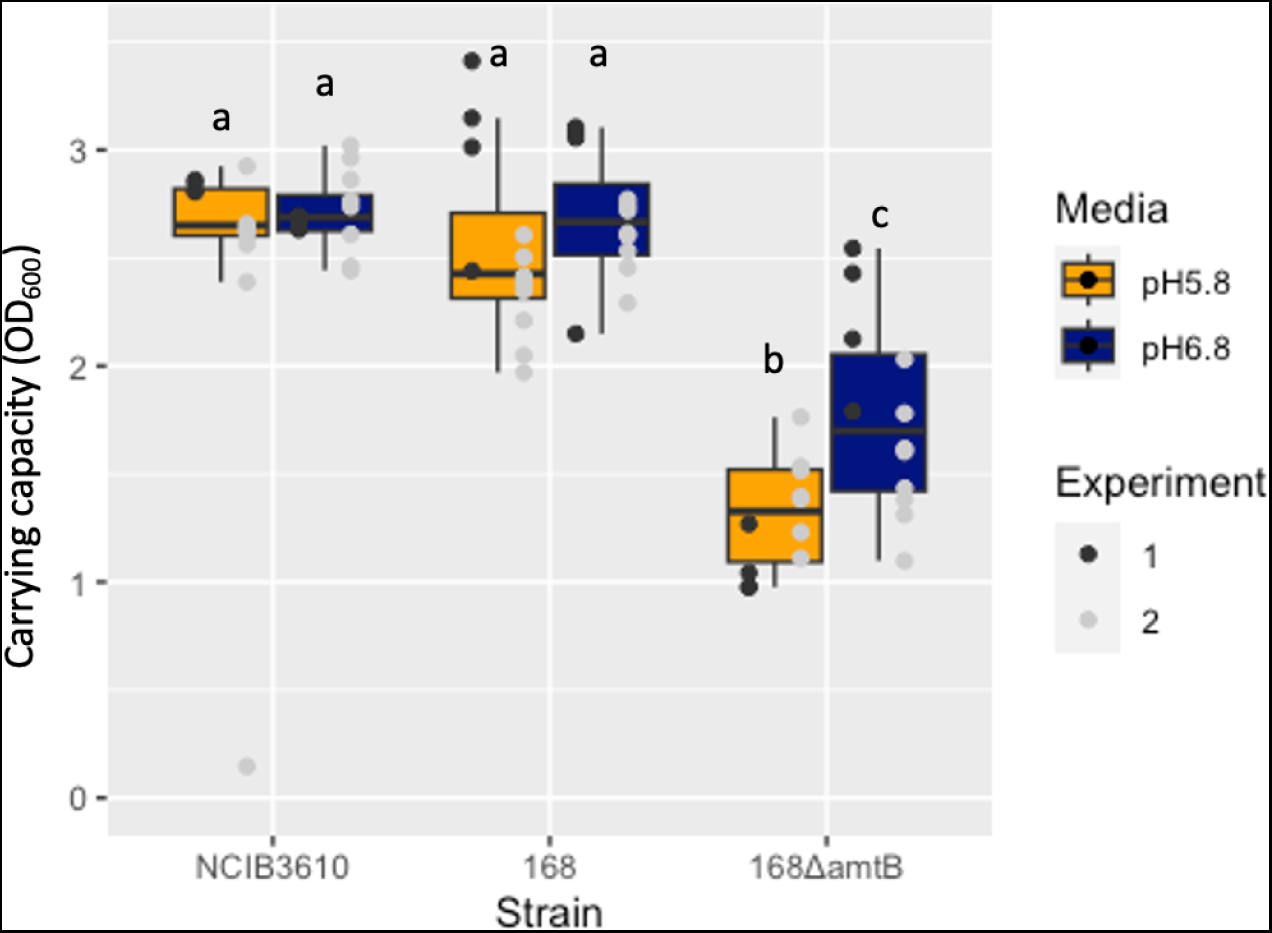
Carrying capacity of fitted bacterial growth curves for wild type *B. subtilis* strains (NCIB3610 and 168) and the ammonium transporter mutant 168ΔamtB in media with two levels of phosphate buffer composition (pH5.8 and pH6.8). Growth curves were measured in a plate reader every 30mins, over the span of 94 hours incubated at 30*◦*C and continually shaken at 200rpm. Experiment was repeated twice, boxes indicate median, LQ and UQ values while the end of whiskers indicate maxima and minima. Individual well measurements are shown with points overlayed on the boxplots. Lettering above boxes indicates significant differences (Pairwise Wilcoxon p*<*0.05)

We also expect the mutant to have higher ammonia leakage (at any pH level). To test this, we have used the wildtype and the mutant strains to initiate co-cultures with *S. indica* on agar plates - we could not use supernatant-based liquid culture experiments, as this *B.subtilis* strain is a tryptophan auxotroph and added amino acid in monoculture would interfere with the ammonium-based interaction. On agar media with only nitrate as nitrogen source (including no tryptophan), we found that both the wildtype and the mutant can enhance *S. indica* growth, but the mutant can do so to a higher degree (Figure S6). This finding supports the idea that the mutant with reduced ammonia uptake results in higher ammonia levels in its environment.

### 2.6 *S. indica* can reduce its external pH and this might impact its ammonoia-based interaction with *B. subtilis*

Another consideration arising from the model is that the spatial distribution of ammonia and ammonium will be dependent on environmental pH and that microorganisms - like *S. indica* - might reduce local pH in order to facilitate ammonium scavenging. Inline with this hypothesis, we found that *S. indica* does indeed reduce the pH of its local environment when grown on agar, whereas *B. subtilis* does not (Figure S9). While such pH reduction could have several causes and consequences, we note here that it would be expected to have an impact on ammonia diffusion around *S.indica* and *B. subtilis*, and might impact their interaction.

To test this hypothesis we repeated our on-plate interaction assay but included repeated addition of potassium phosphate buffer to either stabilise the initial pH (5.8) or bring the pH up over time (pH 6.8) - larger pH alterations were not feasible in our media conditions. We were unable to demonstrate a growth reduction in *S. indica* with the pH stabilisation scenario. This might be due to limitations of our experimental setup, in terms of our ability to control pH and quantify *S. indica* growth accurately. Further experiments with other species and media, where larger pH alterations can be implemented, can further address the possible role of local pH dynamics on ammonium-based interactions.

## 3 Discussion

We have studied here the possibility of nitrogen sharing, among two common, plant-growth promoting microbes, *S. indica* and *B. subtilis*. This specific fungi-bacteria pair is previously shown to present a thiamine-mediated auxotrophic interaction ([Jiang et al., 2018]), and the fungi *S. indica* was shown to be incapable of nitrate assimilation ([Zuccaro et al., 2011]). We re-confirmed the latter proposition and showed that *S. indica* is indeed incapable of growth when nitrate is the sole nitrogen source. We found, however, that this incapacity is lifted, and growth significantly enhanced, in the presence of *B. subtilis*. We find that this effect is mediated through ammonia, which is leaked from *B. subtilis*, when grown on nitrate. We find that these results can be rationalised by a mathematical model, incorporating known permeability of ammonia to cell membranes, active ammonium uptake and assimilation, and ammonia-ammonium equilibrium. Utilising this model, we predict that some level of ammonium leakage is inevitable for cells and becomes worse (more leakage) under low environmental pH and reduced uptake and assimilation rates. Taken together, these results experimentally prove that nitrogen sharing among soil microbes is a feasible and specific interaction mechanism, and that ammonia-based interactions can be influenced by environmental pH around the microbes and their individual ammonia uptake and assimilation rates.

The presented findings are relevant in our understanding of nitrogen dynamics in soils. Nitrate is a major component of global soils ([Hill et al., 2011], ([Lagunas et al., 2023]), [Craswell, 2021], [Walvoord et al., 2003], [Rao and Puttanna, 2000]) and applied fertilisers making it an abundant source of nitrogen for many microorganisms and plants. Assimilating nitrate is an energetically expensive process, necessitating both active uptake of nitrate and its reduction to ammonia ([Moreau et al., 2019], [Lin and Stewart, 1997], [Crawford and Arst Jr, 1993]). Ammonia, free amino acids and more complex organic nitrogen-containing compounds also exist in global soils ([Andresen et al., 2015], [Jones and Darrah, 1994], [Jones and Kielland, 2002], [Warren, 2017]). Microbes and plants have been demonstrated to have to ability to take up not only nitrate and ammonium but also free amino acids and more complex compounds including peptides and proteins ([Hill et al., 2011], [Paungfoo-Lonhienne et al., 2008], [Benjdia et al., 2006], [Ling and Armstead, 1995]). The presented finding that ammonia sharing can be possible for microbes that don’t utilise nitrate suggest that this mechanism can allow such microbes to ‘cut an energetic corner’ in their search for nitrogen and rely on the exuded/ leaked compounds provided by other members of a soil community. This mechanism could then lead to loss of genetic capacity to assimilate nitrate. The high membrane permeability of ammonia means that no organisms are able to hold on to 100% of their reduced ammonium, making a small proportion available for uptake by other organisms.

Interestingly, we have also identified here a small amount of ammonium production in the *S. indica* cultures grown on nitrate media and without bacterial supernatant supplementation. This suggests that *S. indica* can use spore stored nitrogen sources, or recycling of amino acids, to achieve some growth in nitrate media and ammonia is leaked as a consequence. Deamination of nucleic and amino acids, in nitrogen free media, has been observed in response to nitrogen limitation in *Escherichia coli* ([Muse et al., 2003]) and *Neurospora crassa* ([DeBusk and Ogilvie, 1984]) respectively. The observed amounts of ammonium present in *S. indica* supernatants can be achieved given the theoretical amount of nitrogen stored in the spores used. For example, ([Money, 2016]) stated that a 0.4mm diameter fungal spore would weigh approximately 40*µ*g. *S. indica* spores have a diameter approximately x10 less than this value (in the order 10-50*µ*m ([Dias et al., 2020]))and if we assume equal distribution of weight per volume an *S. indica* spore could weigh approximately 40ng. Combining this with the observation that fern spores, of a similar size to *S. indica* spores, have a weight of a similar magnitude ([Gómez-Noguez et al., 2016]), a culture of approximately 50,000 spores per 100mL would give a total of 2mg of spores. Assuming that the total N content of fungal tissue is approximately 5% ([Manzi et al., 1999]), we can expect a total of 7.1*µ*mol nitrogen in 100mL, translating to a nitrogen concentration of 71*µ*M. In our experiments we detect around 50*µ*M NH_4_^+^ in *S. indica*, grown in nitrate only media. Given the estimates we present, that would translate to a 60-70% loss of nitrogen, originally stored in the spore, into the supernatant. Considering that this process of spore-stored nitrogen assimilation and ammonia loss would be happening in tandem with ammonia assimilation from *B. subtilis* supernatant, we conclude that growth of *S. indica* in media supplemented with *B. subtilis* supernatant results in net ammonia consumption.

The findings presented here suggest that interactions among microbes, based on ammonia leakage and consumption, can be influenced by environmental pH. In particular, any nitrogen assimilating microbe will suffer at low pH, leaking more ammonia, while any microbe relying on such leaked ammonia would benefit from lower pH in its local environment, as long as it has high ammonia uptake and assimilation rates. In other words, a reduced capacity to actively uptake ammonium would be a greater disadvantage in a low pH environment. These considerations can be used to speculate that microbes relying solely on ammonia as nitrogen source could use high uptake rates and acidifying their local pH as an effective strategy to scavenge ammonia from their environment. On the other hand, microbes able to use multiple nitrogen sources might not have specifically high ammonium uptake rates, or might suffer from lowered pH more (in terms of leaking more ammonia). Experimental results presented here align with these considerations, in that we found an ammonium transporter mutant of *B. subtilis* to be more susceptible to reduced growth in face of environmental pH reduction, and that *S. indica* promotes a low pH local environment upon growth. Additionally, *B. subtilis* may represent a particularly good “leaker” of ammonia; *B. subtilis* GDH may exclusively function to degrade, rather than produce, glutamate owing to a very low affinity for the ammonium ion ([Commichau et al., 2008], [Gunka and Commichau, 2012]). This would mean that in *B. subtilis* ammonia is only assimilated via GS. This would effectively reduce the assimilation rate described by our model and lead to higher rates of leakage into exterior media.

These results, and pH impacts on nitrogen competition in the environment, is interesting to consider in the soil context. Low pH around Arabidopsis root systems has been shown to be vital in the suppression of plant immune responses by *Pseudomonas* ([Yu et al., 2019a]). Beneficial microbes also have to suppress plant immune responses to colonise host tissue ([Yu et al., 2019b], [Jacobs et al., 2011]). Thus, it is possible that pH lowering is a strategy used by *S. indica*, or other fungi, initially to aid in host immune responses to allow tissue colonisation and has subsequently allowed *S. indica* to utilise low pH environments and high ammonium uptake as an ammonium-scavenging strategy.

## 4 Methods

### 4.1 ATS media

Here we use a modified version of the ATS media described by[Lincoln et al., 1990]. The composition of this media is a follows: 70*µ*M H_3_BO_3_, 14*µ*M MnCl_2_, 0.5*µ*M CuSO_4_, 1*µ*M ZnSO4, 0.2*µ*M NaMoO_4_, 10*µ*M NaCl, 0.01*µ*M CoCl_2_, 2.5mM KPO_4_ (2.3mM KH_2_PO_4_ and 0.2mM K_2_HPO_4_ for pH5.8), 3mM MgSO_4_, 3mM CaCl_2_, 50*µ*M Fe-EDTA, 100mM glucose, 500nM thiamine. N-sources were added in the form of 10mM KNO_3_, 5mM (NH_4_)_2_SO_4_ or 5mM glutamine. For plate experiments 2% agarose was added as a gelling agent.

### 4.2 S. indica propagation

*S. indica* spores were stored at -80*^◦^*C in 0.02% tween_20_ solution at a density of 500,000 spores/mL. Spores were then germinated on CM-agar plates (20g/L glucose, 6g/L NaNO_3_, 2g/L peptone, 1g/L casein hydrolysate, 1g/L yeast extract, 1.52g/L KH_2_PO_4_, 502mg/L MgSO_4_*·*7H_2_O, 502mg/L KCl, 6mg/L MnCl_2_*·*4H_2_O, 2.65mg/L ZnSO_4_*·*H_2_O, 1.5mg/L H_3_BO_3_, 0.75mg/L KI, 0.13mg/L CuSO_4_*·*5H_2_O, 2.4ng/L Na_2_MO_4_*·*2H_2_O, 15g/L agar) and allowed to mature for 4 weeks. Plugs were taken with a cork borer from these “master plates” and placed into the centre of fresh CM-agar plates with mycelia in contact with the fresh media. These primary plates were allowed to mature for a further 8 weeks minimum before spore harvesting. Spores were harvested from these plates by washing with 0.02% tween_20_ solution and adjusting to a concentration of 500,000 spores/mL for use as inoculum in later experiments. Spore density was assessed using a Fuchs-Rosenthal haemocytometer.

### 4.3 B. subtilis propagation

*B. subtilis* strains NCIB3610, 168 and 168ΔamtB were maintained at -80*^◦^*C in 25% glycerol. A scraping from this glycerol was streaked on LB-agar (10g/L Peptone, 5g/L Yeast extract, 10g/L NaCl, 1.5% Agar) plates and incubated at 30*^◦^*C overnight. Single colonies from these plates were inoculated into 10mL LB-broth for overnight cultures. After overnight growth the cells were spun at 12,000 x g for 1min and washed twice with 100mM NaCl and adjusted to the desired optical density (OD_600_ = 0.5 unless otherwise stated) for use as inoculum in later experimental procedures.

### 4.4 Plate inoculation of S.indica and B. subtilis experiments

For solid media experiments 2mL of 2% agarose ATS media in 3.5cm petri dishes was used as described above and N-sources were added at the concentrations indicated. 5*µ*L of *S. indica* spore suspension was inoculated centrally onto plates for mono-culture experiments, plates were sealed with parafilm and allowed to growth for 3 weeks in the dark in static 30*^◦^*C incubator. For co-culture experiments, the *S. indica* inoculum was slightly offset to one side to allow for 2cm separation between organisms. These plates were placed in static 30*^◦^*C incubator for 2 day prior to *B. subtilis* inoculation. After 2 days plates were opened and allowed to dry for around 1 hr in sterile conditions. 1*µ*L of *B. subtilis* inoculum or 1*µ*L 100mM NaCl was placed on the opposite side of the plate. Plates were wrapped in parafilm and kept in the 30*^◦^*C static incubator until imaging. For pH manipulation experiments the plates were unwrapped every 2 days for two weeks and had 50*µ*L 1M KPO_4_, either at pH 5.8 or 6.8, added to the S. indica inoculation site, allowed to dry for approximately 30mins and re-wrapped at returned to the 30*^◦^*C incubator. On days 4 and 6, 50*µ*L of 0.01% bromocresol purple was also added to the centre of one plate per treatment plates to guide further addition of buffer. Beyond day 6, bromocresol was stable so no more bromocresol treatments were included.

### 4.5 Preparation of B. subtilis supernatant

500mL conical flasks were sterilised and filled to a final volume of 200mL ATS media with 10mM KNO_3_. To this, 1mL of *B. subtilis* inoculum (prepared as described above) was added and cultures were placed in 30*^◦^*C shaking 170rpm. For initial experiments considering a range of *B. subtilis* OD_600_ supernatants cultures were sampled at approximately 24, 28, 32 and 52hrs. For later experiments, cultures were sampled at 28hrs and the exact OD_600_ measured. The cultures were centrifuged at x3200g for 10mins followed by vacuum filter sterilisation of the supernatant through Corning PES filters 0.22µ*µ*m for use in later experiments and analysis.

### 4.6 S. indica liquid culture experiments

*S. indica* liquid cultures were grown in 250mL conical flasks filled to a final volume of 100mL ATS with the indicated N-sources, *B. subtilis* supernatant and 100*µ*L spore inoculum. These were placed in shaking 30*^◦^*C incubator 170rpm for 1 week before sampling. On sampling cultures were passed through Miracloth (Merck-Millipore) to filter mycelia from the growth supernatant. Supernatant was collected for metabolite analysis and mycelia were scraped from the Miracloth surface and placed in Eppendorf tubes to dry for weight measurements. Tubes and samples were dried at 70*^◦^*C for 1-2 days and then returned to room temperature for around 24 hours before weighing to equilibrate to ambient humidity.

### 4.7 Nitrate, ammonium and amino acid quantification

Quantification of free amino acids and ammonium, plus total protein content (amino acid content post acid hydrolysis) was conducted externally by Genaxxon using HPLC (high performance liquid chromatography) LC3000 with post-column ninhydrin derivitisation at 125*^◦^*C. For detection of protein-derived amino acids, samples were hydrolysed, prior to HPLC separation, in 6N HCl at 110*^◦^*C for 60 hours.

For detection of nitrate ions we used the Dionex*^TM^* ICS-5000^+^ ion-chomotography system. We used an anionic column with KOH eluent. 2.5*µ*L of sample (filtered through 0.22*µ*m syringe filter) was injected and run at a flow rate of 0.38mL/min for 30 minutes in a constant gradient as follows 0mins 1.5mM KOH, 8mins 1.5mM KOH, 18mins 15mM KOH, 23mins 24mM KOH, 24mins 60mM KOH, 30mins 60mM KOH. Nitrate was detected at an elution time of around 12.8 minutes using a conductivity detector. Standard curve for quantification is presented in Figure S10B.

### 4.8 Fluorescence pH quantification

Fluorescence standard curve was constructed by preparing KPO_4_ buffer at 4 different pH’s by combining different ratios of 1M KH_2_PO_4_:K_2_HPO_4_ as follows: pH4 100:0 (adjusted down to pH 4 with 0.1M HCl), pH5.8 91.5:8.5, pH6.8 50.3:49.7, pH8 6:94. Propidium iodide (PI) and the pH sensitive 2’,7’-Bis-(2-Carboxyethyl)-5-(and-6)-Carboxyfluorescein (BCECF) (Invitrogen) were added to these at a final concentration of 100*µ*M and 10*µ*M respectively. PI was to act as a normalisation standard to account for general light intensity shifts. Solutions were placed in 3.5cm petri dishes and imaged for fluorescence readings in multiple plate locations. Plates were imaged with bright field illumination with 0.5ms exposure, HcRED1 filter set 41043 (Chroma) exposure (100ms) and EGFP filter set 41018 (Chroma) exposure (100ms). Light source was pE-300white (CoolLED). Standard curve for pH quantification is presented in Figure S10A.

### 4.9 Mathematical modelling

We used a modified version of the model developed by [Kim et al., 2012] to consider the dynamics of ammonia and ammonium between the interior and exterior of a cell growing in a nitrate-only and given pH environment. We defined differential equations for the external (Equation 1) and internal (Equation 2) ammonium concentrations.

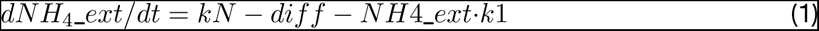

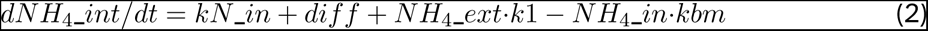

Parameters *kN* and *kN_*in represent the external influx (from cells into the external environment) and internal influx (from nitrate reduction) of ammonium. Parameter *diff* represents diffusion into or out of the cell from the exterior environment and is given by Equation 3 where *PV_A_*is the rate of ammonia diffusion from the cell and *k_i_n* and *k_e_xt* are the internal and external dissociation constants respectively for the ammonium-ammonia equilibrium at the pH of those compartments. The rates *k1* and *kbm* represent the rates of ammonium import into the cell and ammonium incorporation into biomass respectively. For our *kN_i_n* parameter we inferred an amount of ammonium per hour that must be generated through nitrate reduction. We considered in our experiment that there was a reduction of approximately 2.5mM nitrate over the course of the 28hr growth (Figure S5). Thereby indicating at least 90*µ*M per hour (2500/28) of internal ammonium production at the population level.

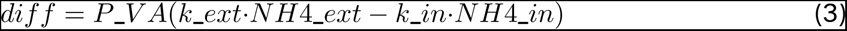

We solved differential equations for change in external ammonium concentration (Equation 1) and internal ammonium concentration (Equation 2) until simulations reached steady state. We check steady state has been reach by verifying that the external ammonium concentration is not changing by more than 10*^−^*^5^ We used a time frame of 25 hours to mimic our experimental set up and the simulated systems had reached or come close to steady state in this time. Endpoint measurements were then used to generate surface plots.

Python code for these simulations is available on github at https://github.com/lukeZrich/RichardsNsharing2024

### 4.10 Plate reader assessment of bacterial growth

Liquid ATS media, as described previously, was used for plate reader growth experiments with the following modifications. KPO_4_ buffer composition was altered to increase the pH by increasing the ratio of K_2_HPO_4_ to KH_2_PO_4_ such that the final media contained 1.3mM KH_2_PO_4_ and 1.2mM K_2_HPO_4_ for pH6.8. With standard ATS composition, increasing the pH causes magnesium and calcium phosphate precipitates to form. To combat this we reduced the concentration of MgSO_4_ and CaCl_2_ by ten fold for 0.3mM each. *B. subtilis* inocula were prepared as described above and 1*µ*L used to inculate 200*µ*L of media per well. Cultures were grown in plate reader (CLARIOstar BMG Labtech) for 3 days at 30*^◦^*C, 200rpm and OD_600_ measurements were taken every 30 minutes.

### 4.11 BLAST searching of nitrogen assimilation genes homology

Protein gene accession as described in Data file S1 were used to perform protein-protein BLAST against the *S. indica* and *S. vermifera* genetic information stored in the NCBI (TaxID 1109443 and 109899 respectively). Default settings were used to run blastp ([Altschul et al., 1997]) protein-protein blast.

## Acknowledgements

We would like to thank Prof. Munehiro Asally at the University of Warwick for providing all of the *B. subtilis* strains presented in this work. We would also like to thank Dr. Kelsey Cremin and Dr. Jerko Rosko for their contribution in helping with imaging microbial growth plates. We dedicate this manuscript to the memory of our beloved Xue Jiang, who made the initial observations of nitrogen sharing studied in this work.

---

**Figure S1:**
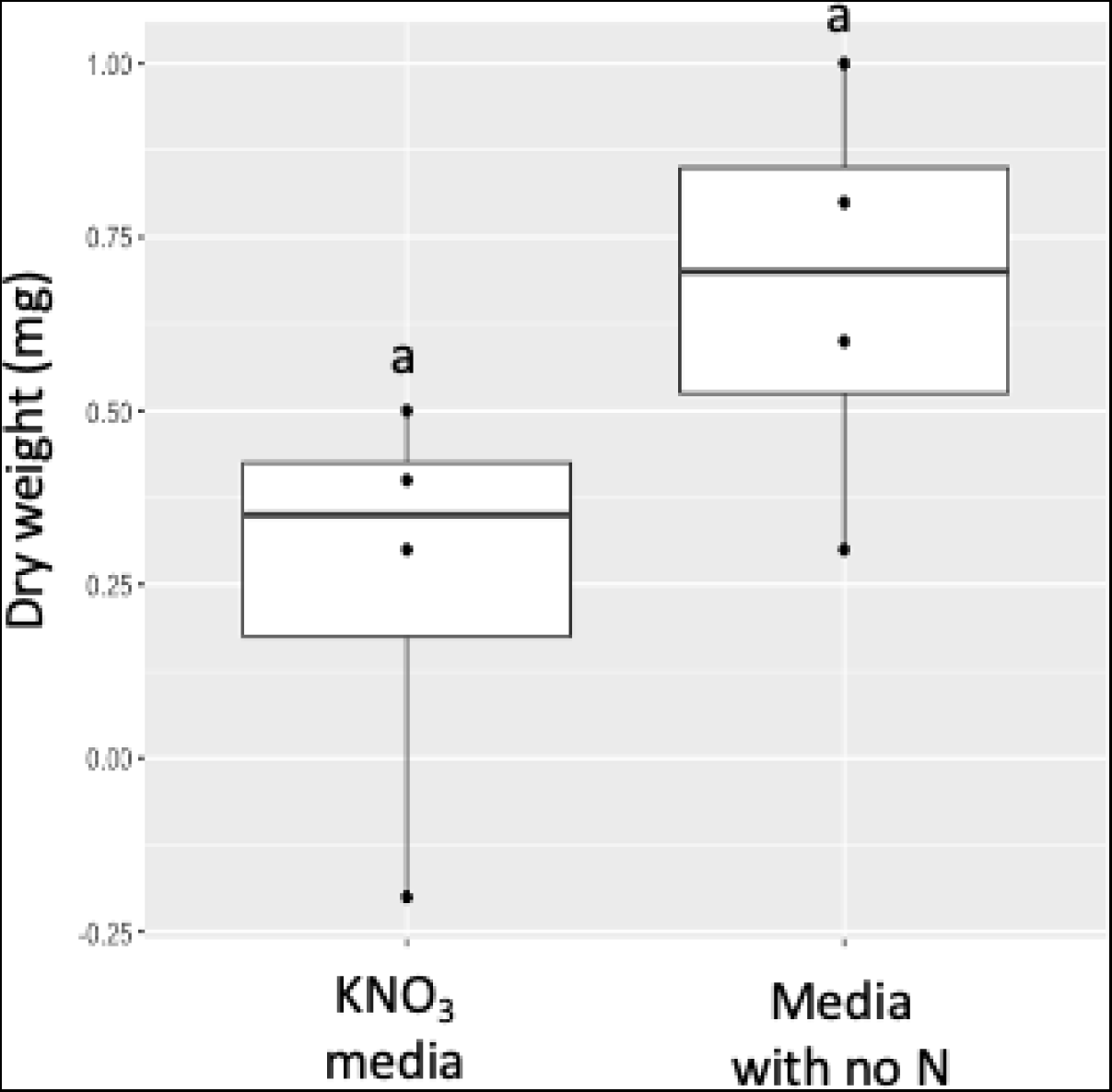
Dry weight of *S.indica* growth after 1 week of liquid culture in ATS media supplemented with 10mM KNO_3_ or no N-source.

**Figure S2:**
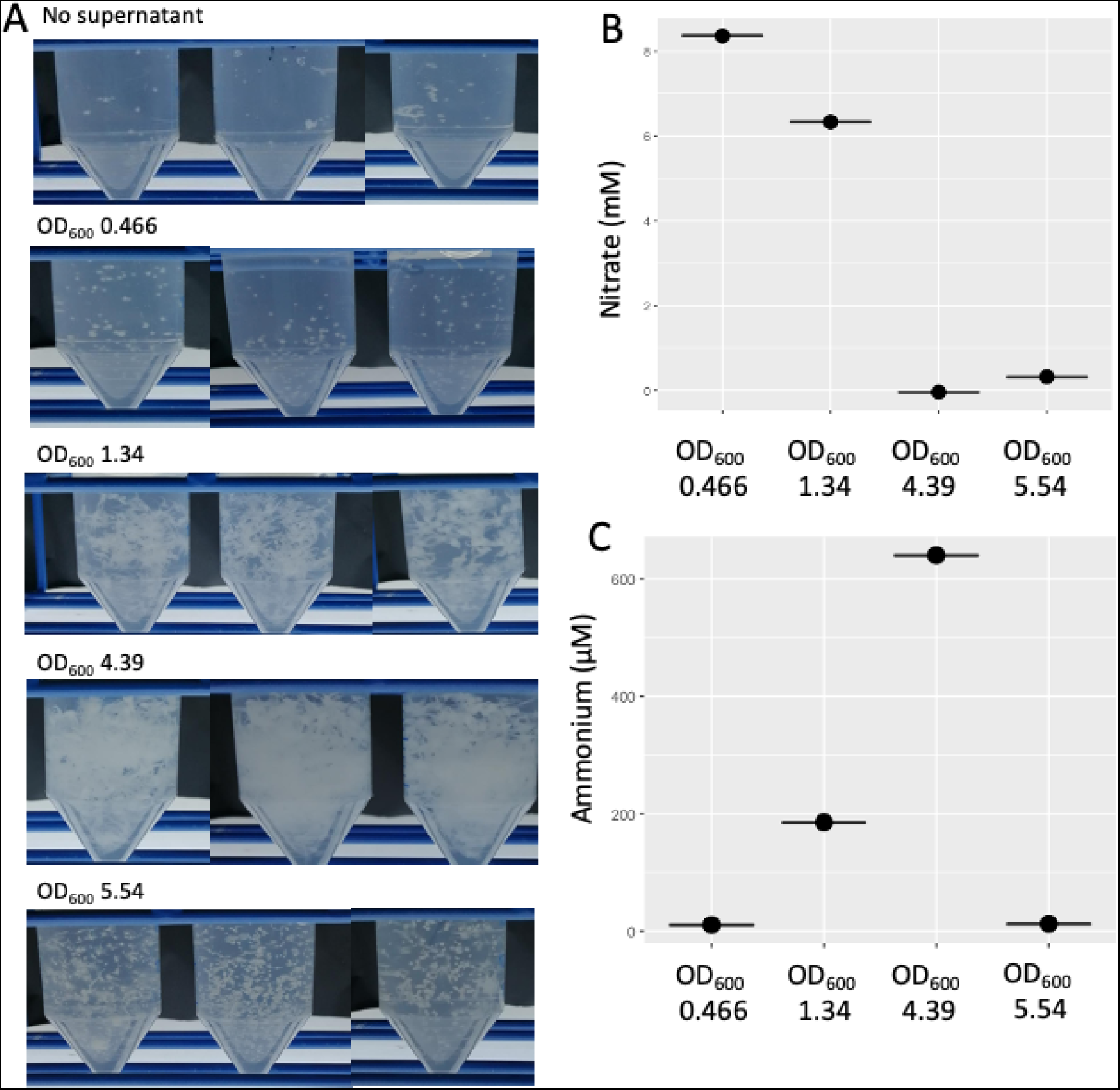
*B. subtilis* growth stage impacts *S. indica* growth]A) *S. indica* growth after 1 week of liquid culture in 50:50 mixtures of ATS 10mM KNO_3_ and *B. subtilis* supernatants grown to different optical densities in the same media. B) Nitrate and C) ammonium quantification for the same *B. subtilis* supernatants used in A.

**Figure S3:**
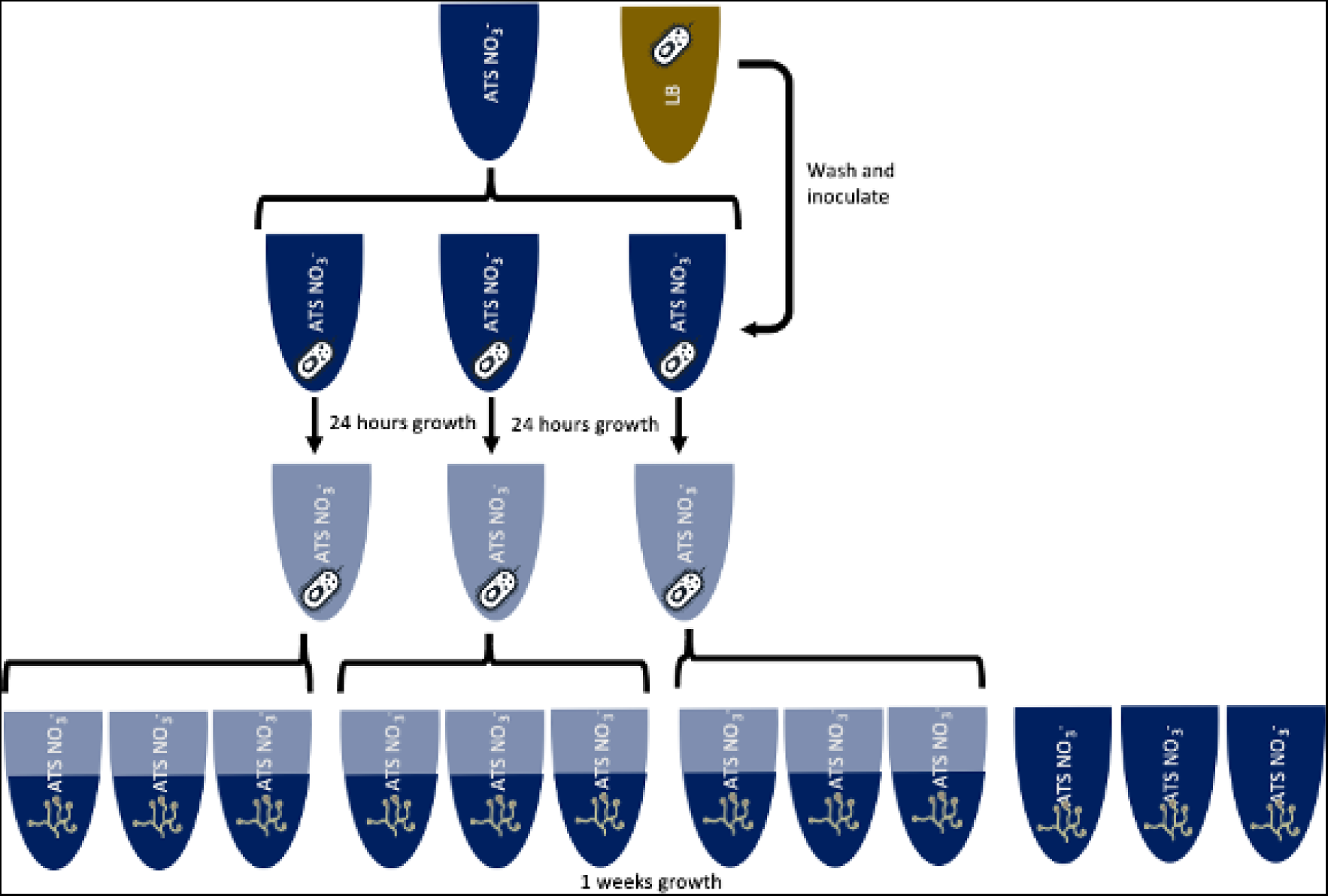
Experimental schematic to explain the basic experimental design for *S. indica* growth quantification in *B. subtilis* supernatants.

**Figure S4:**
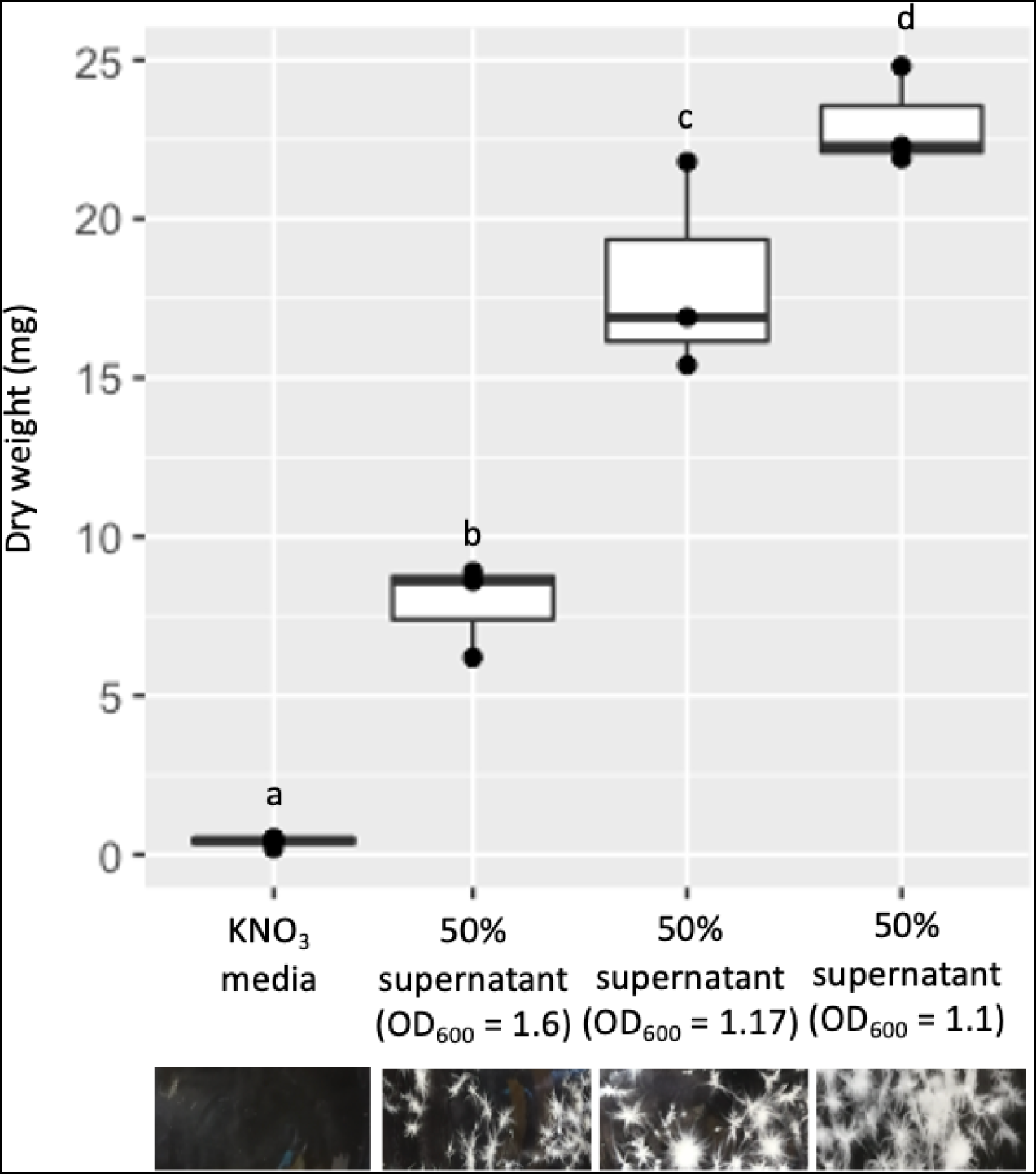
A) Dry weight of *S.indica* growth after 1 week of liquid culture in ATS media supplemented with 10mM KNO_3_ or a 50:50 mixture of KNO_3_-only media and *B. subtilis* supernatant generated from KNO_3_-only media type to a bacterial optical density indicated on the x-axis. Mid point indicates median, edges of boxes indicate lower and upper quartiles and ends of whiskers indicate maxima and minima. Significant differences are indicated by letters above bars (p*<*0.05 Tukey HSD).

**Figure S5:**
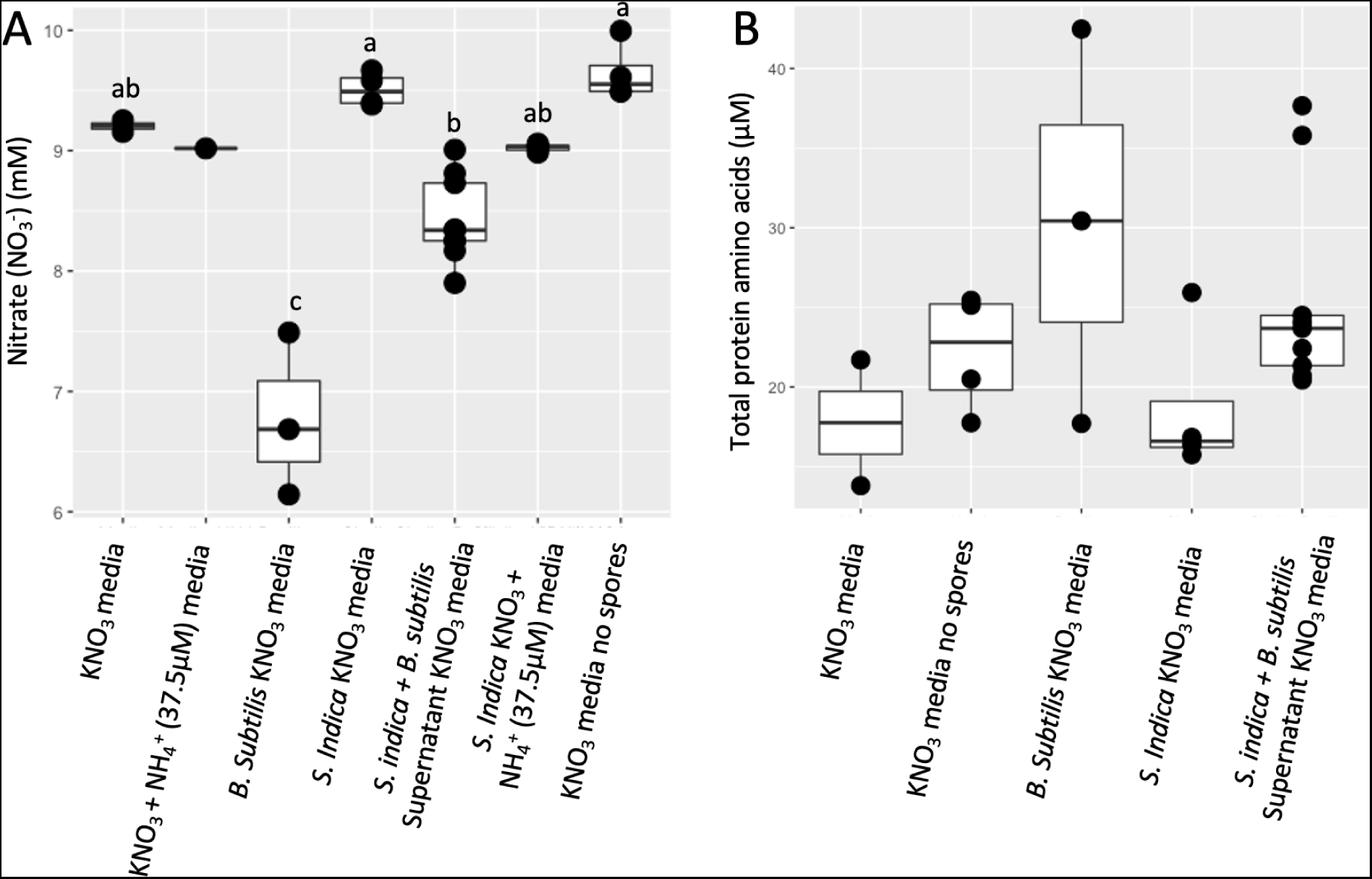
A) Nitrate and B) total protein-derived (after hydrolysis) amino acid quantification for *S. indica* and *B. subtilis* growth supernatants in various media and media only controls as indicated on the x axes. A) Significant differences are indicated with letters above bars (TukeyHSD p*<*0.05) (KNO_3_ + NH_4_^+^ (37.5*µ*M) media was excluded from statistical analysis because this represents only one measurement). B) No statistical differences were found between any of the treatments (TukeyHSD p*<*0.05).

**Figure S6:**
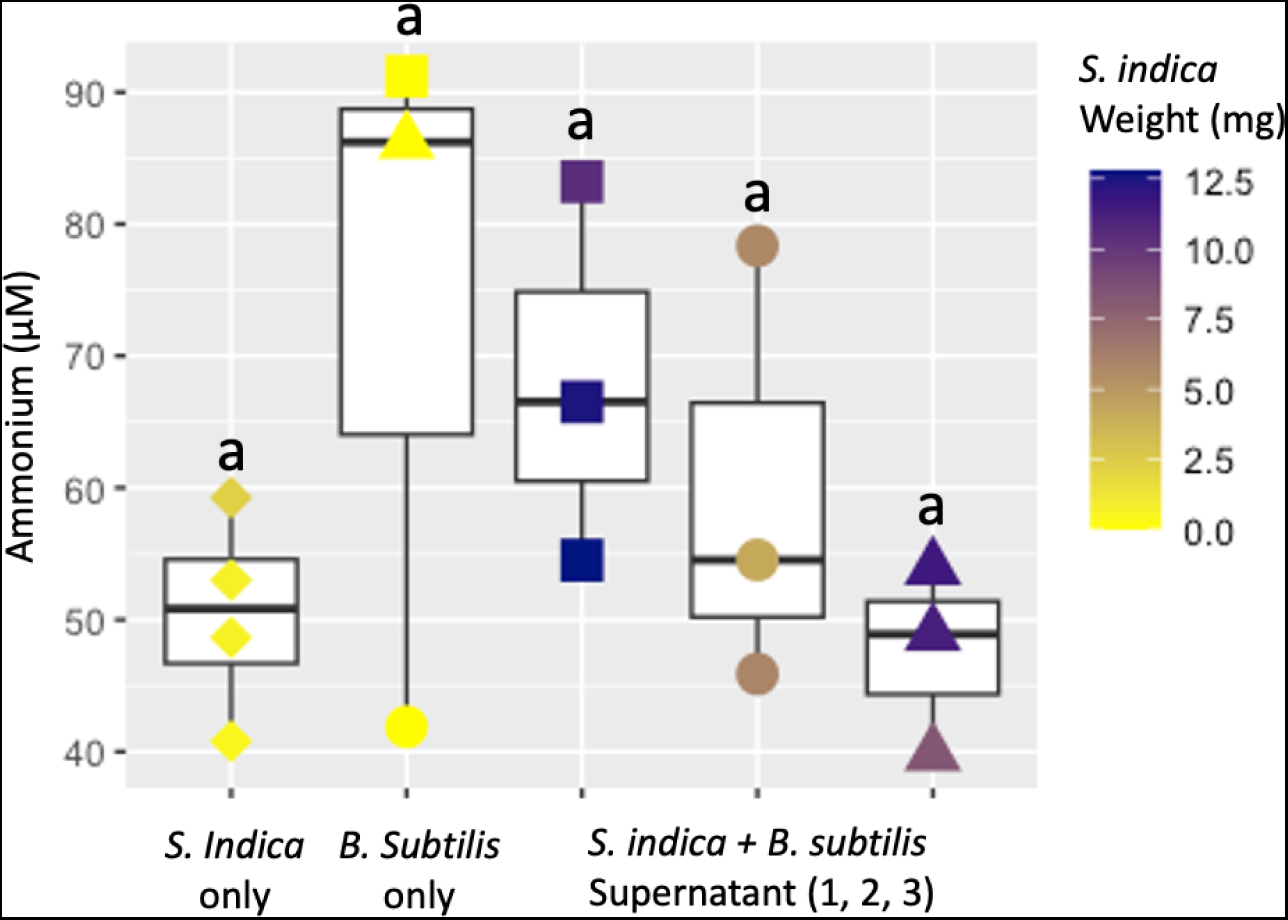
NH_4_^+^ concentration as measured by HPLC in *S. indica* and *B. subtilis* cultures in conditions indicated on the x-axis. Where applicable *S. indica* dry weight (from Figure 3A) for each individual point is indicated with a colour scale. Mid point indicates median, edges of boxes indicate lower and upper quartiles and ends of whiskers indicate maxima and minima. Significant differences are indicated with letters above boxes (TukeyHSD p *<* 0.05). Plot point shapes indicate the OD_600_ of the *B. subtilis* culture at the time of supernatant harvest (from Figure 3A): diamond - NA, square - 1.02, circle - 1.83, triangle - 1.26.

**Figure S7:**
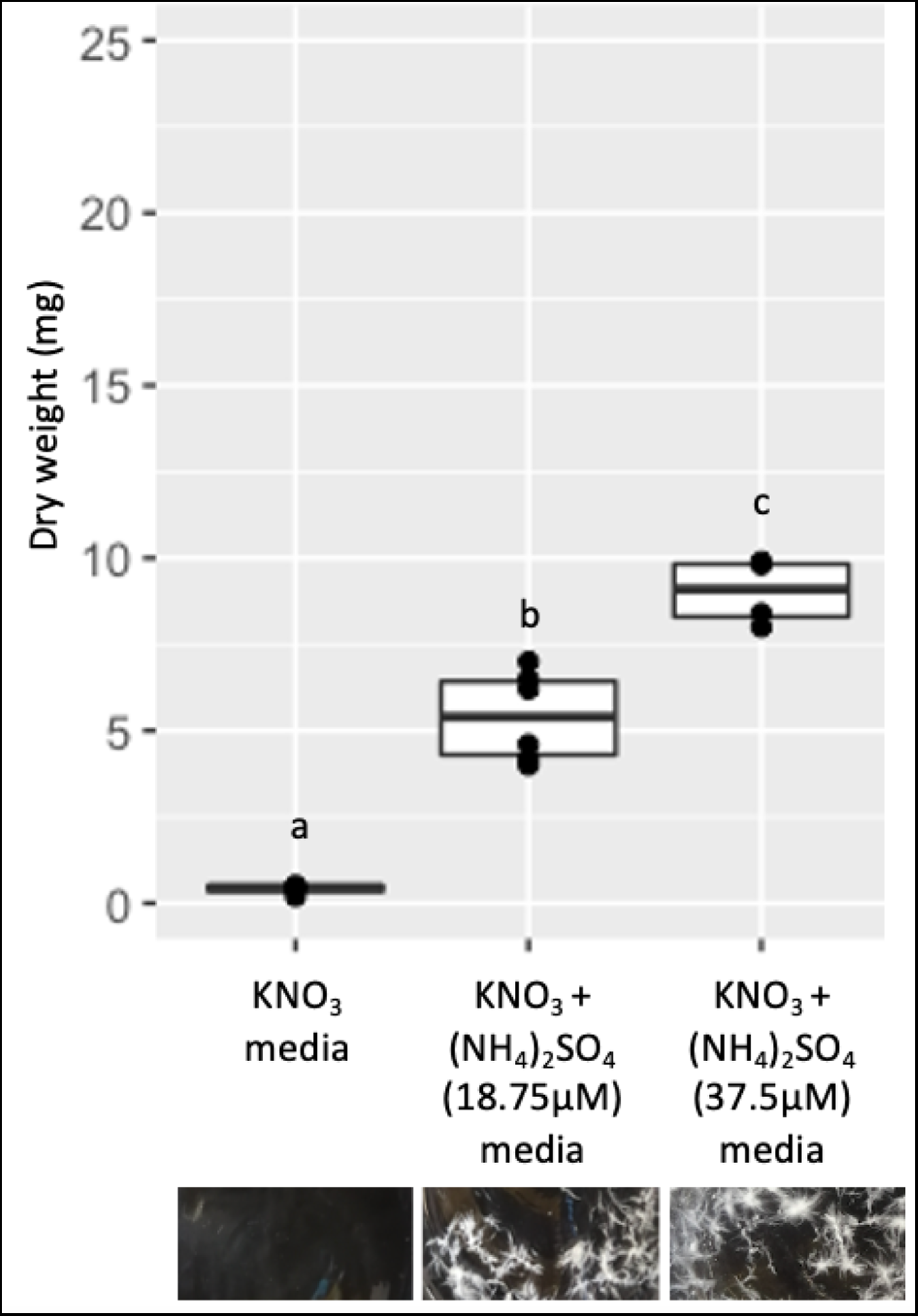
A) Dry weight of *S.indica* growth after 1 week of liquid culture in ATS media supplemented with 10mM KNO_3_, 10mM KNO_3_ + 18.75µM (NH_3_)_2_SO_4_ or 10mM KNO_3_ + 37.5µM (NH_3_)_2_SO_4_. Mid point indicates median, edges of boxes indicate lower and upper quartiles and ends of whiskers indicate maxima and minima. Significant differences are indicated by letters above bars (p*<*0.05 Tukey HSD).

**Figure S8:**
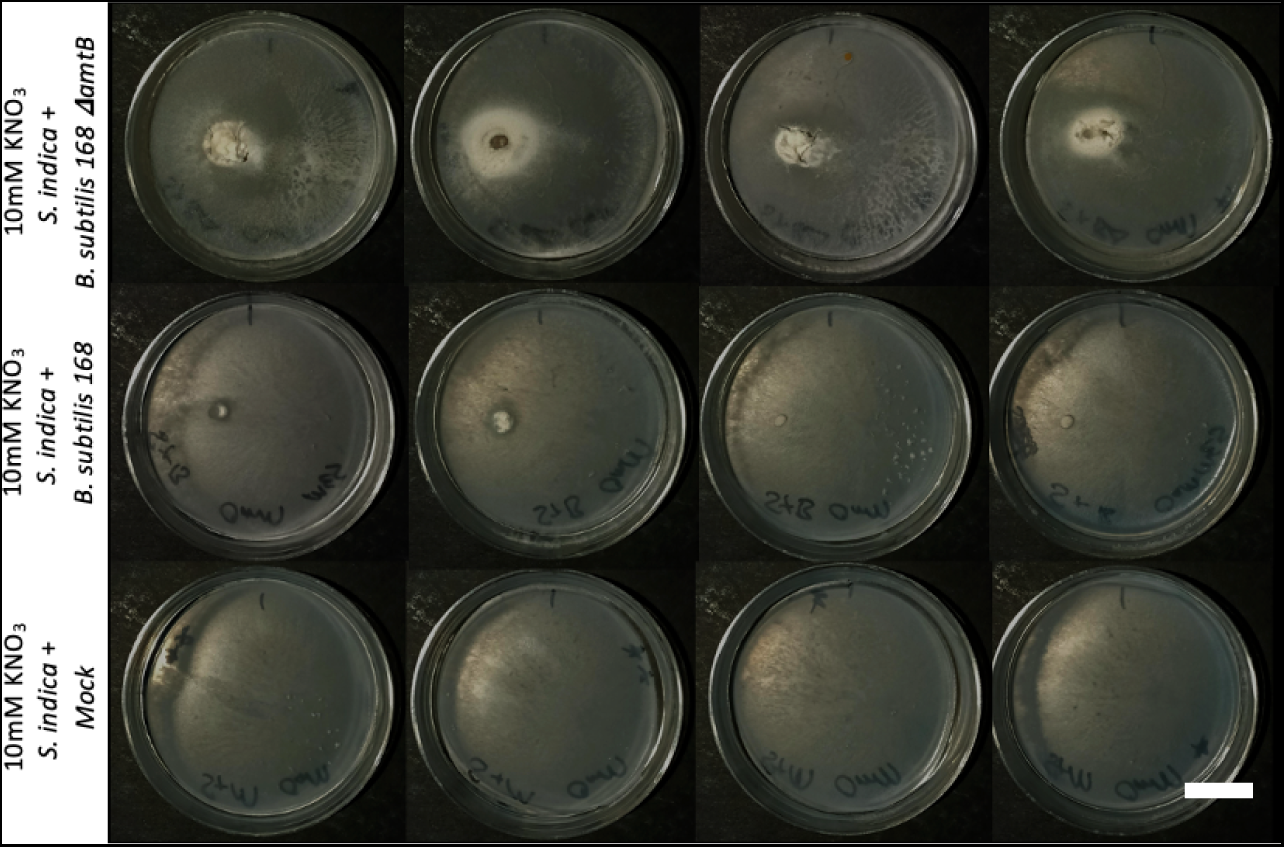
*S. indica* grown in isolation or in the presence of *B. subtilis* 168 or the ammonium uptake mutant on ATS media supplemented with 10mM KNO_3_ for 22 days. *B. subtilis* inoculum was added on the RHS of plates 2 days after *S. indica* inoculation on the LHS. On co-culture plates *B. subtilis* colonies are not visible at the site of inoculation. In co-culture the *S. indica* mycelia appears more dense at the site of inoculation. Scale bar is 1cm.

**Figure S9:**
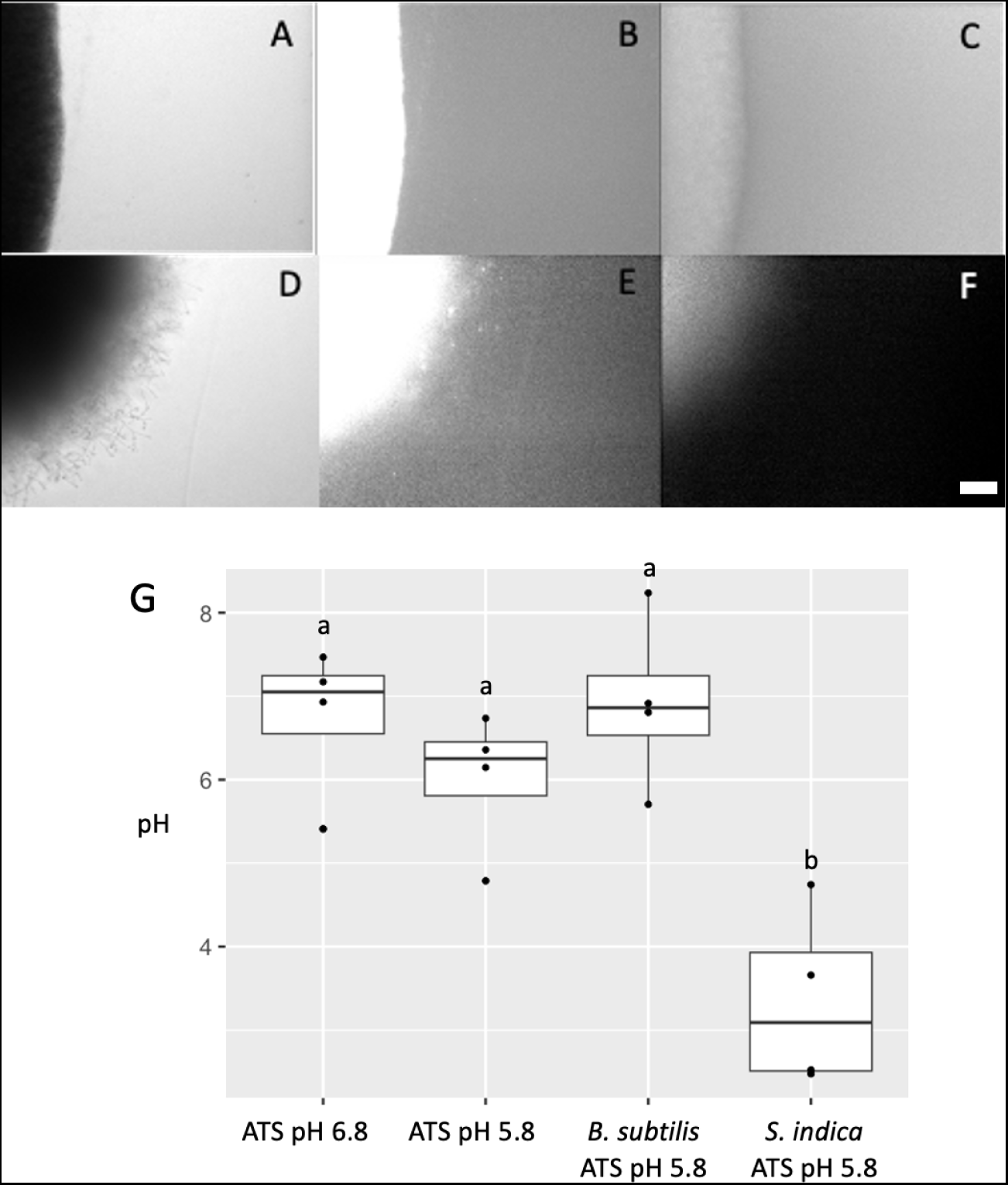
A and D) Bright field, B and E) RED fluorescence (PI) and, C and F) GFP fluorescence (BCECF pH sensitive) images of A, B and C) *B. subtilis* and, D, E and F) *S. indica* grown in ATS media pH5.8 and flooded with 10*µ*M BCECF pH sensitive dye and 100µM PI (propidium iodide). Scale bar indicates 100*µ*m. G shows pH values calculated using BCECF fluorescence values normalised against RED fluorescence for two abiotic media plates at pH 5.8 and 6.8, and *B. subtilis* and *S. indica* plates grown at pH 5.8. Fluorescence images for pH calculations were taken in an are directly adjacent to grown colonies but without colonies in view.

**Figure S10:**
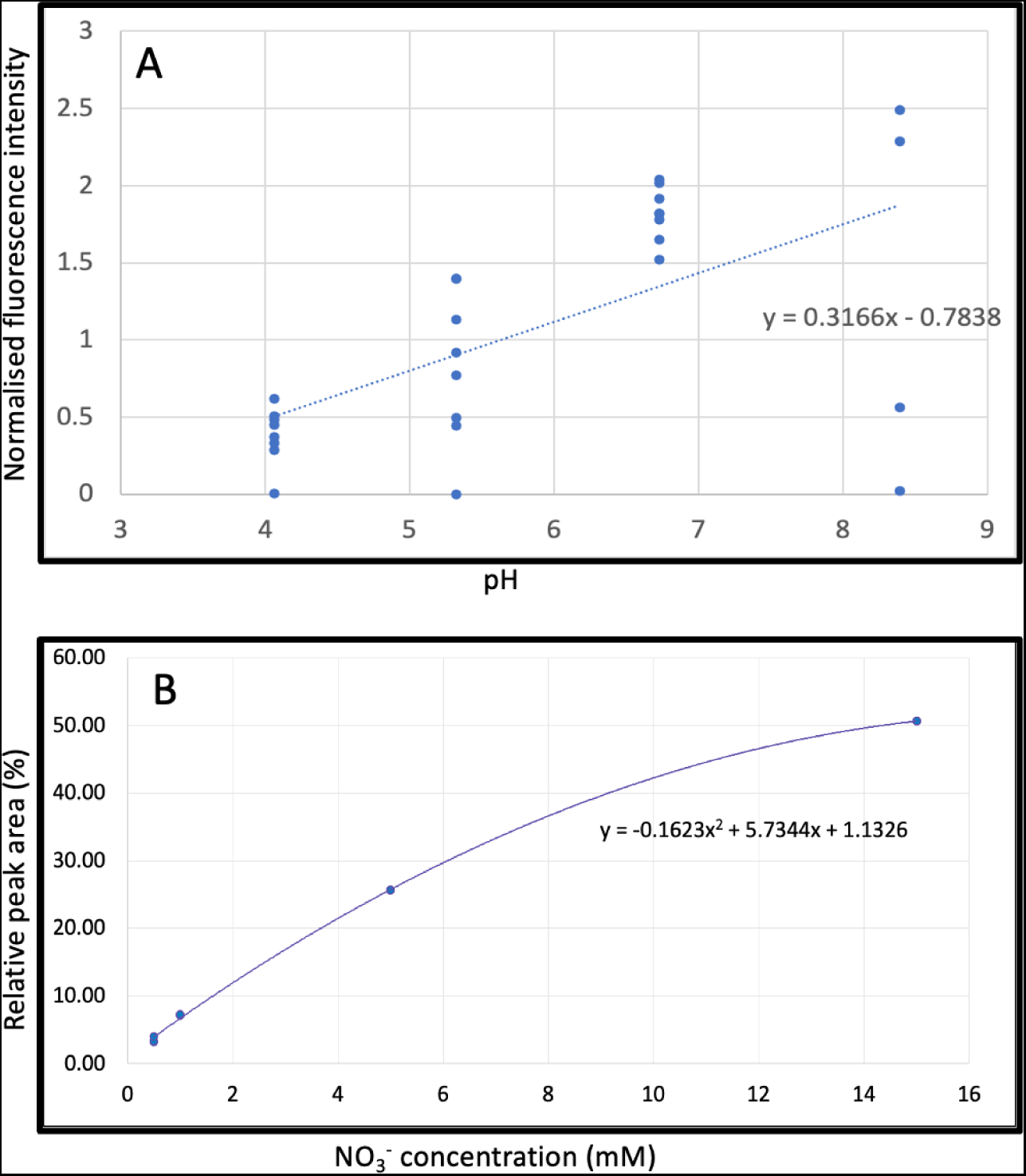
Standard curves and associated equations used for the quantification of A) pH and B) nitrate.

## References

Altschul, S. F., Madden, T. L., Schäffer, A. A., Zhang, J., Zhang, Z., Miller, W., and Lipman, D. J. (1997). Gapped blast and psi-blast: a new generation of protein database search programs. Nucleic acids research, 25(17):3389–3402.

Andresen, L., Bodé, S., Tietema, A., Boeckx, P., and Rütting, T. (2015). Amino acid and n mineralization dynamics in heathland soil after long-term warming and repetitive drought. Soil, 1(1):341–349.

ÁVILA, J., GONZÁLEZ, C., Brito, N., and Siverio, J. M. (1998). Clustering of the yna1 gene encoding a zn (ii) 2cys6 transcriptional factor in the yeast hansenula polymorpha with the nitrate assimilation genes ynt1, yni1 and ynr1, and its involvement in their transcriptional activation. Biochemical journal, 335(3):647–652.

Bakker, P. A., Berendsen, R. L., Van Pelt, J. A., Vismans, G., Yu, K., Li, E., Van Bentum, S., Poppeliers, S. W., Gil, J. J. S., Zhang, H., et al. (2020). The soil-borne identity and microbiome-assisted agriculture: looking back to the future. Molecular plant, 13(10):1394–1401.

Barea, J.-M. and Richardson, A. E. (2015). Phosphate mobilisation by soil microorganisms. Principles of Plant-Microbe Interactions: Microbes for Sustainable Agriculture, pages 225–234.

Benjdia, M., Rikirsch, E., Müller, T., Morel, M., Corratgé, C., Zimmermann, S., Chalot, M., Frommer, W. B., and Wipf, D. (2006). Peptide uptake in the ectomycorrhizal fungus hebeloma cylindrosporum: characterization of two di-and tripeptide transporters (hcptr2a and b). New Phytologist, 170(2):401–410.

Bhattacharjya, S., Adhikari, T., Kundu, S., Sahu, A., and Patra, A. K. (2019). Evaluation of microbial solubilisation of nano rock phosphate. Int J Curr Microbiol App Sci, 8(01):1055–1069.

Campbell, W. H. and Kinghorn, J. R. (1990). Functional domains of assimilatory nitrate reductases and nitrite reductases. Trends in biochemical sciences, 15(8):315–319.

Cavaliere, M., Feng, S., Soyer, O. S., and Jiménez, J. I. (2017). Cooperation in microbial communities and their biotechnological applications. Environmental microbiology, 19(8):2949–2963.

Commichau, F. M., Gunka, K., Landmann, J. J., and Stülke, J. (2008). Glutamate metabolism in bacillus subtilis: gene expression and enzyme activities evolved to avoid futile cycles and to allow rapid responses to perturbations of the system. Journal of bacteriology, 190(10):3557–3564.

Condron, L., Stark, C., O’Callaghan, M., Clinton, P., and Huang, Z. (2010). The role of microbial communities in the formation and decomposition of soil organic matter. Soil microbiology and sustainable crop production, pages 81–118.

Coyte, K. Z. and Rakoff-Nahoum, S. (2019). Understanding competition and cooperation within the mammalian gut microbiome. Current Biology, 29(11):R538–R544.

Craswell, E. (2021). Fertilizers and nitrate pollution of surface and ground water: an increasingly pervasive global problem. SN Applied Sciences, 3(4):518.

Crawford, N. M. and Arst Jr, H. N. (1993). The molecular genetics of nitrate assimilation in fungi and plants. Annual review of genetics, 27(1):115–146.

DeBusk, R. M. and Ogilvie, S. (1984). Participation of an extracellular deaminase in amino acid utilization by neurospora crassa. Journal of bacteriology, 159(2):583–589.

Dias, T., Pimentel, V., Cogo, A. J. D., Costa, R., Bertolazi, A. A., Miranda, C., de Souza, S. B., Melo, J., Carolino, M., Varma, A., et al. (2020). The free-living stage growth conditions of the endophytic fungus serendipita indica may regulate its potential as plant growth promoting microbe. Frontiers in Microbiology, 11:562238.

D’Souza, G. and Kost, C. (2016). Experimental evolution of metabolic dependency in bacteria. PLoS genetics, 12(11):e1006364.

Eichmann, R., Richards, L., and Schäfer, P. (2021). Hormones as go-betweens in plant microbiome assembly. The Plant Journal, 105(2):518–541.

Fu, H., Uchimiya, M., Gore, J., and Moran, M. A. (2020). Ecological drivers of bacterial community assembly in synthetic phycospheres. Proceedings of the National Academy of Sciences, 117(7):3656–3662.

Gómez-Noguez, F., Pérez-García, B., Mehltreter, K., Orozco-Segovia, A., and Rosas-Pérez, I. (2016). Spore mass and morphometry of some fern species. Flora, 223:99–105.

Gourmelon, V., Maggia, L., Powell, J. R., Gigante, S., Hortal, S., Gueunier, C., Letellier, K., and Carriconde, F. (2016). Environmental and geographical factors structure soil microbial diversity in new caledonian ultramafic substrates: a metagenomic approach. Plos one, 11(12):e0167405.

Gunka, K. and Commichau, F. M. (2012). Control of glutamate homeostasis in bacillus subtilis: a complex interplay between ammonium assimilation, glutamate biosynthesis and degradation. Molecular microbiology, 85(2):213–224.

Hill, P. W., Quilliam, R. S., DeLuca, T. H., Farrar, J., Farrell, M., Roberts, P., Newsham, K. K., Hopkins, D. W., Bardgett, R. D., and Jones, D. L. (2011). Acquisition and assimi-lation of nitrogen as peptide-bound and d-enantiomers of amino acids by wheat. PLoS One, 6(4):e19220.

Hoek, T. A., Axelrod, K., Biancalani, T., Yurtsev, E. A., Liu, J., and Gore, J. (2016). Resource availability modulates the cooperative and competitive nature of a microbial cross-feeding mutualism. PLoS biology, 14(8):e1002540.

Jacobs, S., Zechmann, B., Molitor, A., Trujillo, M., Petutschnig, E., Lipka, V., Kogel, K.-H., and Schäfer, P. (2011). Broad-spectrum suppression of innate immunity is required for colonization of arabidopsis roots by the fungus piriformospora indica. Plant physiology, 156(2):726–740.

Jiang, X., Zerfaß, C., Feng, S., Eichmann, R., Asally, M., Schäfer, P., and Soyer, O. S. (2018). Impact of spatial organization on a novel auxotrophic interaction among soil microbes. The ISME journal, 12(6):1443–1456.

Jones, D. L. and Darrah, P. (1994). Amino-acid influx at the soil-root interface of zea mays l. and its implications in the rhizosphere. Plant and soil, 163:1–12.

Jones, D. L. and Kielland, K. (2002). Soil amino acid turnover dominates the nitrogen flux in permafrost-dominated taiga forest soils. Soil biology and biochemistry, 34(2):209–219.

Kazamia, E., Czesnick, H., Nguyen, T. T. V., Croft, M. T., Sherwood, E., Sasso, S., Hodson, S. J., Warren, M. J., and Smith, A. G. (2012). Mutualistic interactions between vitamin b12-dependent algae and heterotrophic bacteria exhibit regulation. Environmental microbiology, 14(6):1466–1476.

Kehe, J., Ortiz, A., Kulesa, A., Gore, J., Blainey, P. C., and Friedman, J. (2021). Positive interactions are common among culturable bacteria. Science advances, 7(45):eabi7159.

Kim, M. and Or, D. (2017). Hydration status and diurnal trophic interactions shape microbial community function in desert biocrusts. Biogeosciences, 14(23):5403–5424.

Kim, M., Zhang, Z., Okano, H., Yan, D., Groisman, A., and Hwa, T. (2012). Need-based activation of ammonium uptake in escherichia coli. Molecular systems biology, 8(1):616.

Kraemer, S. (2004). Iron oxide dissolution and solubility in the presence of siderophores. Aquatic sciences, 66(1):3–18.

Kuypers, M. M., Marchant, H. K., and Kartal, B. (2018). The microbial nitrogen-cycling network. Nature Reviews Microbiology, 16(5):263–276.

Lagunas, B., Richards, L., Sergaki, C., Burgess, J., Pardal, A. J., Hussain, R. M., Richmond, B. L., Baxter, L., Roy, P., Pakidi, A., et al. (2023). Rhizobial nitrogen fixation efficiency shapes endosphere bacterial communities and medicago truncatula host growth. Microbiome, 11(1):146.

Lauber, C. L., Hamady, M., Knight, R., and Fierer, N. (2009). Pyrosequencing-based assessment of soil ph as a predictor of soil bacterial community structure at the continental scale. Applied and environmental microbiology, 75(15):5111–5120.

Lemanceau, P., Bauer, P., Kraemer, S., and Briat, J.-F. (2009). Iron dynamics in the rhizosphere as a case study for analyzing interactions between soils, plants and microbes.

Lin, J. T. and Stewart, V. (1997). Nitrate assimilation by bacteria. Advances in microbial physiology, 39:1–30.

Lincoln, C., Britton, J. H., and Estelle, M. (1990). Growth and development of the axr1 mutants of arabidopsis. The Plant Cell, 2(11):1071–1080.

Ling, J. and Armstead, I. (1995). The in vitro uptake and metabolism of peptides and amino acids by five species of rumen bacteria. Journal of Applied Bacteriology, 78(2):116–124.

Manzi, P., Gambelli, L., Marconi, S., Vivanti, V., and Pizzoferrato, L. (1999). Nutrients in edible mushrooms: an inter-species comparative study. Food chemistry, 65(4):477–482.

Money, N. P. (2016). Spore production, discharge, and dispersal. In The fungi, pages 67–97. Elsevier.

Moreau, D., Bardgett, R. D., Finlay, R. D., Jones, D. L., and Philippot, L. (2019). A plant perspective on nitrogen cycling in the rhizosphere. Functional Ecology, 33(4):540–552.

Muse, W. B., Rosario, C. J., and Bender, R. A. (2003). Nitrogen regulation of the codba (cytosine deaminase) operon from escherichia coli by the nitrogen assimilation control protein, nac. Journal of bacteriology, 185(9):2920–2926.

Osborne, R., Rehneke, L., Lehmann, S., Roberts, J., Altmann, M., Altmann, S., Zhang, Y., Köpff, E., Dominguez-Ferreras, A., Okechukwu, E., et al. (2023). Symbiont-host interactome mapping reveals effector-targeted modulation of hormone networks and activation of growth promotion. Nature Communications, 14(1):4065.

Paungfoo-Lonhienne, C., Lonhienne, T. G., Rentsch, D., Robinson, N., Christie, M., Webb, R. I., Gamage, H. K., Carroll, B. J., Schenk, P. M., and Schmidt, S. (2008). Plants can use protein as a nitrogen source without assistance from other organisms. Proceedings of the National Academy of Sciences, 105(11):4524–4529.

Ponomarova, O., Gabrielli, N., Sévin, D. C., Mülleder, M., Zirngibl, K., Bulyha, K., Andrejev, S., Kafkia, E., Typas, A., Sauer, U., et al. (2017). Yeast creates a niche for symbiotic lactic acid bacteria through nitrogen overflow. Cell systems, 5(4):345–357.

Qiang, X., Weiss, M., Kogel, K.-H., and Schäfer, P. (2012). Piriformospora indica—a mutualistic basidiomycete with an exceptionally large plant host range. Molecular plant pathology, 13(5):508–518.

Rao, E. P. and Puttanna, K. (2000). Nitrates, agriculture and environment. Current Science, 79(9):1163–1168.

Ratzke, C., Barrere, J., and Gore, J. (2020). Strength of species interactions determines biodiversity and stability in microbial communities. Nature ecology & evolution, 4(3):376–383.

Santoyo, G., Pacheco, C. H., Salmerón, J. H., and León, R. H. (2017). The role of abiotic factors modulating the plant-microbe-soil interactions: toward sustainable agriculture. a review. Spanish journal of agricultural research, 15(1):13.

Siverio, J. M. (2002). Assimilation of nitrate by yeasts. FEMS Microbiology Reviews, 26(3):277–284.

Uehling, J. K., Entler, M. R., Meredith, H. R., Millet, L. J., Timm, C. M., Aufrecht, J. A., Bonito, G. M., Engle, N. L., Labbé, J. L., Doktycz, M. J., et al. (2019). Microflu-idics and metabolomics reveal symbiotic bacterial–fungal interactions between mortierella elongata and burkholderia include metabolite exchange. Frontiers in microbiology, 10:2163.

Varma, A., Verma, S., Sudha, Sahay, N., Bütehorn, B., and Franken, P. (1999). Piriformospora indica, a cultivable plant-growth-promoting root endophyte. Applied and environmental Microbiology, 65(6):2741–2744.

Walvoord, M. A., Phillips, F. M., Stonestrom, D. A., Evans, R. D., Hartsough, P. C., Newman, B. D., and Striegl, R. G. (2003). A reservoir of nitrate beneath desert soils. Science, 302(5647):1021–1024.

Warren, C. R. (2017). Variation in small organic n compounds and amino acid enantiomers along an altitudinal gradient. Soil Biology and Biochemistry, 115:197–212.

Weiß, M., Waller, F., Zuccaro, A., and Selosse, M.-A. (2016). Sebacinales–one thousand and one interactions with land plants. New Phytologist, 211(1):20–40.

Yu, K., Liu, Y., Tichelaar, R., Savant, N., Lagendijk, E., van Kuijk, S. J., Stringlis, I. A., van Dijken, A. J., Pieterse, C. M., Bakker, P. A., et al. (2019a). Rhizosphere-associated pseudomonas suppress local root immune responses by gluconic acid-mediated lowering of environmental ph. Current Biology, 29(22):3913–3920.

Yu, K., Pieterse, C. M., Bakker, P. A., and Berendsen, R. L. (2019b). Beneficial microbes going underground of root immunity. Plant, Cell & Environment, 42(10):2860–2870.

Zuccaro, A., Lahrmann, U., Güldener, U., Langen, G., Pfiffi, S., Biedenkopf, D., Wong, P., Samans, B., Grimm, C., Basiewicz, M., et al. (2011). Endophytic life strategies decoded by genome and transcriptome analyses of the mutualistic root symbiont piriformospora indica. PLoS pathogens, 7(10):e1002290.

